# Dissecting components of the *Campylobacter jejuni fetMP-fetABCDEF* gene cluster in iron scavenging

**DOI:** 10.1101/2023.07.05.547857

**Authors:** Tomas Richardson-Sanchez, Anson C. K. Chan, Brendil Sabatino, Helen Lin, Erin C. Gaynor, Michael E. P. Murphy

## Abstract

*Campylobacter jejuni* is a leading cause of bacterial gastroenteritis worldwide. Acute infection can be antecedent to highly debilitating long-term sequelae. Expression of iron acquisition systems is vital for *C. jejuni* to survive the low iron availability within the human gut. The *C. jejuni fetMP-fetABCDEF* gene cluster is known to be upregulated during human infection and under iron limitation. While FetM and FetP have been functionally linked to iron transport in prior work, here we assess the contribution by each of the downstream genes (*fetABCDEF*) to *C. jejuni* growth during both iron-depleted and iron-replete conditions. Significant growth impairment was observed upon disruption of *fetA*, *fetB, fetC*, and *fetD*, suggesting a role in iron acquisition for each encoded protein. FetA expression was modulated by iron-availability but not dependent on the presence of FetB, FetC, FetD, FetE or FetF. Functions of the putative thioredoxins FetE and FetF were redundant in iron scavenging, requiring a double deletion (Δ*fetEF*) to exhibit a growth defect. *C. jejuni* FetE was expressed and the structure solved to 1.50 Å, revealing structural similarity to thiol-disulfide oxidases. Functional characterization in biochemical assays showed that FetE reduced insulin at a slower rate than *E. coli* Trx and that together, FetEF promoted substrate oxidation in cell extracts, suggesting that FetE (and presumably FetF) are oxidoreductases that can mediate oxidation *in vivo*. This study advances our understanding of the contributions by the *fetMP-fetABCDEF* gene cluster to virulence at a genetic and functional level, providing foundational knowledge towards mitigating *C. jejuni*-related morbidity and mortality.

## IMPORTANCE

*Campylobacter jejuni* is a bacterium that is prevalent in the ceca of farmed poultry such as chickens. Consumption of ill-prepared poultry is thus the most common route by which *C. jejuni* infects the human gut to cause a typically self-limiting but severe gastrointestinal illness that can be fatal to very young, old, or immunocompromised people. The lack of a vaccine and an increasing resistance to current antibiotics highlight a need to better understand the mechanisms that make *C. jejuni* a successful human pathogen. This study focused on the functional components of one such mechanism – a molecular system that helps *C. jejuni* thrive despite the restriction on growth-available iron by the human body which typically defends against pathogens. In providing a deeper understanding of how this system functions, this study contributes towards the goal of reducing the enormous global socioeconomic burden caused by *C. jejuni*.

## INTRODUCTION

*Campylobacter jejuni* is a Gram-negative ε-proteobacterium and a leading cause of bacterial diarrheal disease worldwide. *C. jejuni* is a commensal in the intestinal mucosa of many wild and domesticated animals, commonly colonizing farmed poultry flocks, and then undergoing zoonotic transmission to humans, by for example consumption of undercooked poultry or cross-contamination of other food with raw poultry juice (1, 2). Acute human infection results in severe watery to bloody diarrhea and can also be antecedent to highly debilitating long-term sequelae such as inflammatory bowel diseases and autoimmune disorders (3–5). The high socioeconomic burden and impact on human health has been made worse by the inability to produce an effective vaccine and the increasing levels of antibiotic resistance in isolates from both hospitals (6, 7) and poultry meat (8).

Successful *C. jejuni* colonization of the human intestinal mucosa is dependent on a range of factors, including the expression of systems to acquire essential micronutrients such as iron. Recent transcriptional studies on *C. jejuni* have demonstrated upregulation of an eight gene cluster (*CJJ81176_1649-1656,* hereinafter named *fetMP-fetABCDEF*), during human infection (9) and, as our groups have shown, upon exposure to human fecal metabolites (10), indicating a likely role in pathogenesis. The two upstream genes, *fetM* (*CJJ81176_1649*) and *fetP* (*CJJ81176_1650,* also known as *p19*), encode the FetMP iron transport system, which has been shown to be important for growth under iron-limited conditions (11, 12). The six downstream genes *fetABCDEF* (*CJJ81176_1651-1656*) have not previously been characterized individually, but collectively have been shown to be strongly upregulated alongside *fetMP* during iron restriction and upon deletion of *fur*, which encodes the ferric uptake regulator protein (13–15). Two other studies have observed *C. jejuni* growth defects under iron restriction upon disruption of the *fetMP-fetABCDEF* gene cluster, either by deletion of *fetP* (11) or, as our groups recently demonstrated, simultaneous deletion of all six downstream genes (Δ*fetABCDEF*) (10), with growth defects restored upon iron supplementation or complementation with the wild-type gene cluster. Additionally, *C. jejuni* exhibits a biphasic phenotype to the antibiotic streptomycin, where the usual concentration-dependent inhibitory effect is interrupted by a span of streptomycin-tolerant growth at moderate antibiotic concentrations. This tolerance is lost upon deletion of *fetABCDEF* but can be restored with iron supplementation. *C. jejuni* Δ*fetABCDEF* also has increased acid sensitivity and higher resistance to oxidative stress (10). These transcriptional and phenotypic studies link *fetABCDEF* to *C. jejuni* pathogenesis and iron acquisition, providing greater impetus to investigate each individual component of the *fetMP-fetABCDEF* gene cluster.

Gene clusters homologous to *fetMP-fetABCDEF* have been identified in 33 diverse bacterial species across 6 phyla, including 21 species that are associated with human disease (10). The *C. jejuni fetMP-fetABCDEF* gene cluster spans a genomic region of 8.1 kb and consists of two upstream genes (*fetMP*) separated from six downstream genes (*fetABCDEF*) by an 82 base intergenic region (**Figure 1A**). Upstream of the *fetM* start codon is a Fur binding sequence (10 bases upstream) and a putative primary transcription start site (54 bases upstream) (16), suggesting *fetMP-fetABCDEF* may be transcribed as one operon.

**Figure 1:**
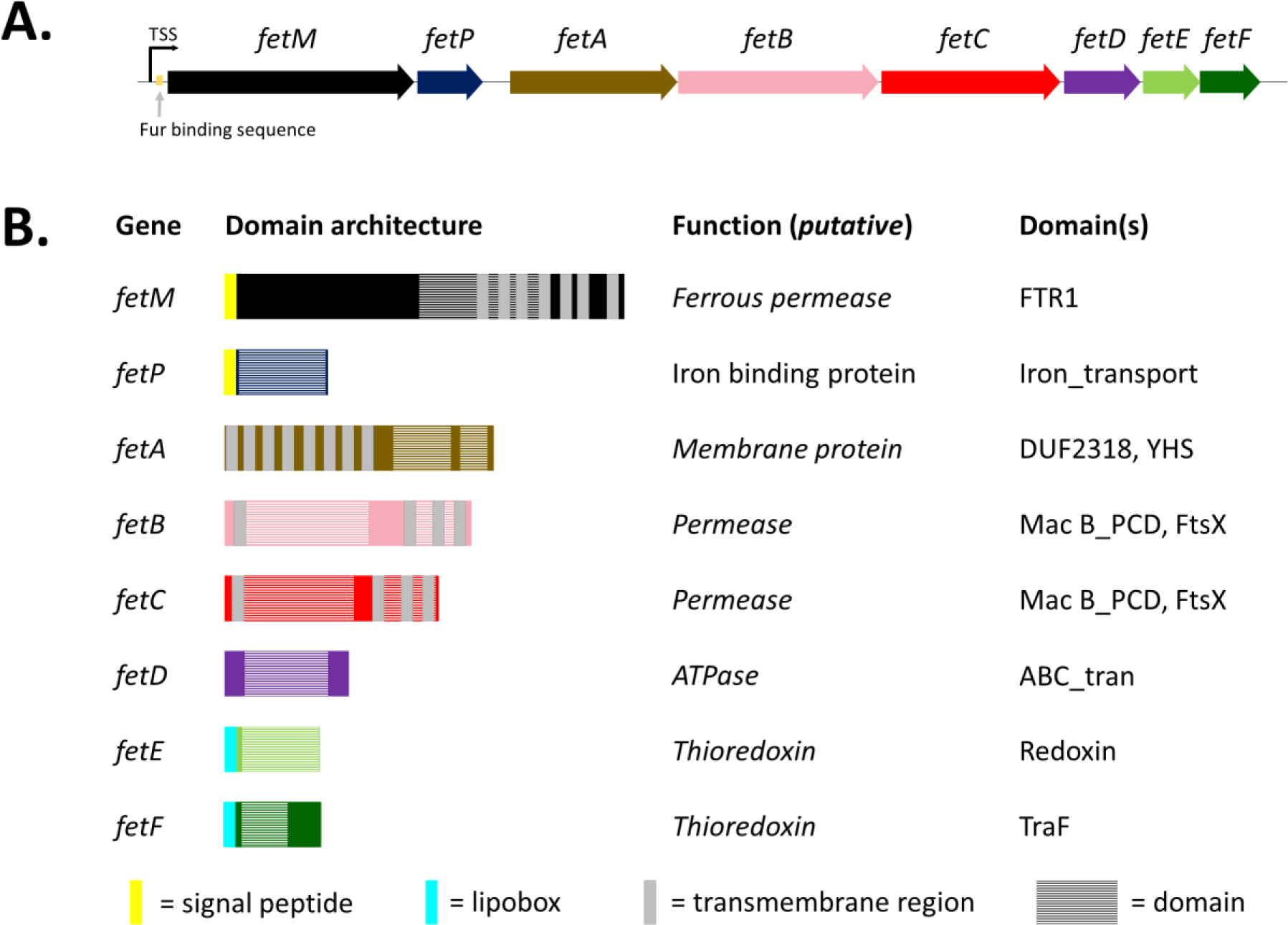
Operonic structure, domain architecture, and model with predicted functions of *fetMP-fetABCDEF* gene cluster (**A**) Proposed operon structure in *C. jejuni* wild-type strain 81-176 with primary transcription start site (TSS) and Fur binding sequence. (**B**) Gene function and domain architecture. Signal peptides and protein lipidation sites (lipoboxes) predicted by SignalP-5.0, transmembrane helices by TMHMM Server v2.0, and domain predictions by Pfam 32.0.

*In silico* domain analysis of the proteins encoded by *fetMP-fetABCDEF* (**Figure 1B**) allows identification of putative functions in cases where functional studies on the proteins or their homologues are lacking. FetM is yet to be characterized in *C. jejuni*, although the *Escherichia coli* homologue has been identified as an iron permease of the oxidase-dependent iron transporter family (17, 18). FetP has been characterized as a periplasmic iron binding protein in *C. jejuni* (11), as well as in *E. coli* (18), *Bordetella* (19), and *Y. pestis* (20). The FetMP iron uptake system was suggested to import ferric-rhodotorulic acid (unpublished data, Stintzi *et al*., 2008) (21), but studies have demonstrated that *C. jejuni* cannot utilise this siderophore for growth (22, 23).

No prior functional studies have been reported for FetABCDEF or their homologues. We predict that these six gene products include a putative membrane permease (encoded by *fetA*), a putative ATP-binding cassette (ABC) transporter (encoded by *fetBCD*), and two putative thioredoxins (encoded by *fetEF*). FetA is predicted to contain eight transmembrane domains that may form a transport channel, a domain of unknown function (DUF2318), and a YHS domain. The genes *fetB* and *fetC* encode domains consistent with ABC transporter permeases, and *fetD* encodes conserved sequence motifs that are vital for ATP binding and hydrolysis. Thus, *fetBCD* is predicted to encode an ABC transporter for active transport of substrates across the inner membrane. Overall, the *fetMP-fetABCDEF* gene cluster is predicted to encode three distinct inner membrane transporters, which would be unusual for an iron uptake system. The genes *fetE* and *fetF* are predicted to encode single domain, membrane-associated thioredoxin oxidoreductases. Oxidoreductases can mediate transitions between Cys-Cys disulfide and dithiol groups within proteins, a function dependent on a conserved active site motif (CXXC) (24) that is present in the FetE (<u>CPSC</u>) and FetF (<u>CGVC</u>) protein sequences.

To ascribe the phenotypes observed for Δ*fetABCDEF* to specific genes within the cluster and to provide insight into the essentiality of each functional unit for iron acquisition, this study used a genetic approach to test individual *fet* gene deletion strains for sensitivity to iron availability and to the antibiotic streptomycin. Comparable degrees of growth impairment were observed under low iron upon disruption of *fetM*, *fetP*, *fetA*, *fetB, fetC*, and *fetD*. Based on single and double mutant analyses, we predict that *fetE* and *fetF* encode gene products that functioned redundantly in iron scavenging. Sensitivity to iron restriction generally corresponds to an increased sensitivity to streptomycin, implicating iron homeostasis as a determinant of growth modality during streptomycin exposure. A molecular biology approach was then used to demonstrate that FetA expression is iron-regulated, and structural biology and biochemical assays allowed further investigation into the function of FetE as a thiol-disulfide oxidoreductase that may act as an oxidase *in vivo*.

## RESULTS

### Sensitivity to iron availability for *C. jejuni* strains

To investigate the contribution of the *fetMP-fetABCDEF* gene cluster components towards growth under different levels of iron availability, gene deletion (Δ) and complemented (^C^) *C. jejuni* strains were constructed for *fetM*, *fetA*, *fetB*, *fetC*, *fetD*, *fetE*, and *fetF* (**Supplemental Figure S1**). To account for potential redundancy of the putative thioredoxins FetE and FetF, a double deletion (Δ*fetEF*) and corresponding complemented (*fetEF*^C^) strain were also constructed. Wild-type *C. jejuni* 81-176 was used as a control, and *C. jejuni* strains corresponding to *fetP* (Δ*fetP*, *fetP*^C^) and the six gene *fet* cluster (Δ*fetABCDEF*, *fetABCDEF*^C^) were used as standards in growth curve experiments (10, 11).

All *C. jejuni* strains were cultured in iron-restricted Mueller-Hinton (MH) broth, standard MH broth, and iron-supplemented MH broth (**Figures 2 and 3, Supplemental Figures S2 and S3**). Iron restriction was achieved by supplementing MH broth with the high affinity ferric iron chelator desferrioxamine B (DFO). Iron supplementation was achieved by supplementing MH broth with 100 µM ferric chloride. As the iron content in MH medium varies between brands and product batches (25), all growth experiments used a single batch of MH medium and the DFO concentration was optimized to 5 µM from a test range of 0 – 20 µM (data not shown). The total Fe content of the standard growth medium was measured at 7 µM by inductively coupled plasma mass spectrometry (ICP-MS). Growth of *C. jejuni* strains under different levels of iron availability was monitored by OD_600_ (**Figure 2**) and colony forming units (CFU, **Figure 3**). Sensitivity of individual strains to changes in iron availability was assessed by comparing cell densities (OD_600_) after 30 h of growth under each condition (**Supplemental Figure S4**).

**Figure 2:**
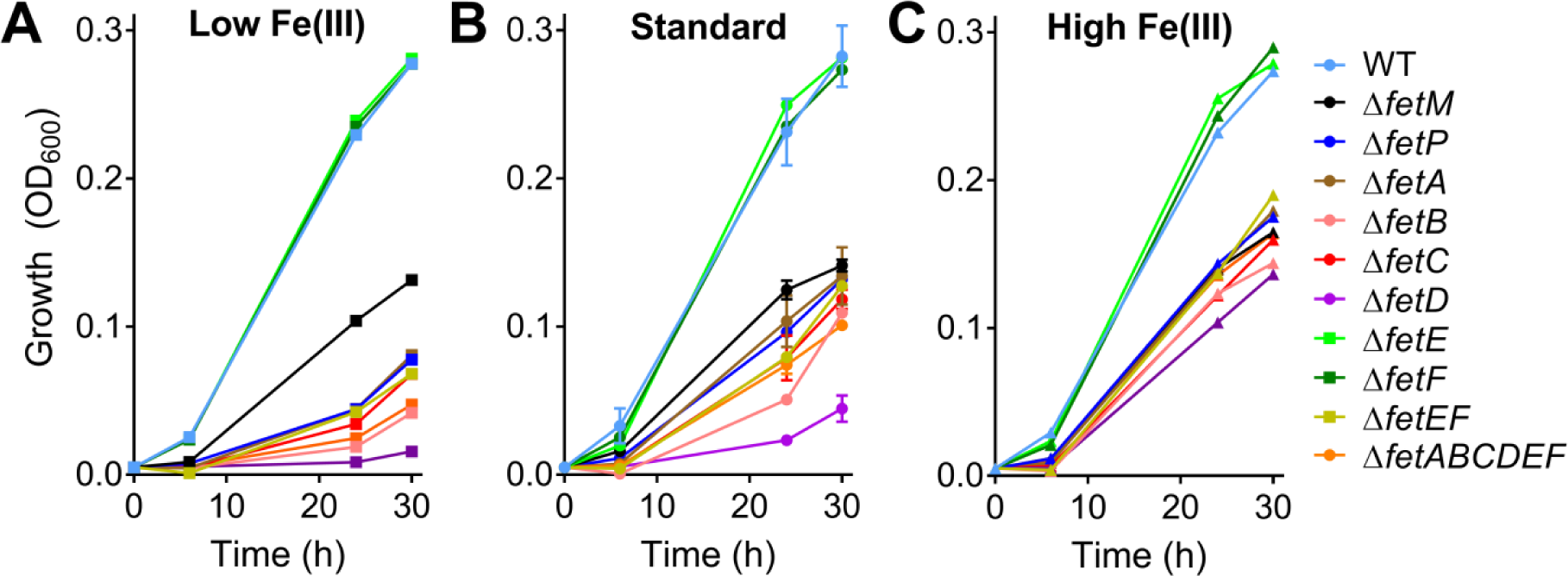
Growth by monitoring OD_600_ for *C. jejuni* gene deletion strains under different levels of iron availability. *C. jejuni* strains were cultured under low (MH + 5 µM DFO, **A**), standard (MH, **B**), or high (MH + 100 µM FeCl_3_, **C**) Fe(III) availability, with growth monitored by absorbance at 600 nm. Corresponding results for *C. jejuni* complemented strains are in **Supplemental Figure S2**. Each strain was assayed in triplicate except for wild-type (WT), which was cultured for every growth experiment and hence was assayed with 18 biological replicates. Error bars represent standard deviation.

**Figure 3:**
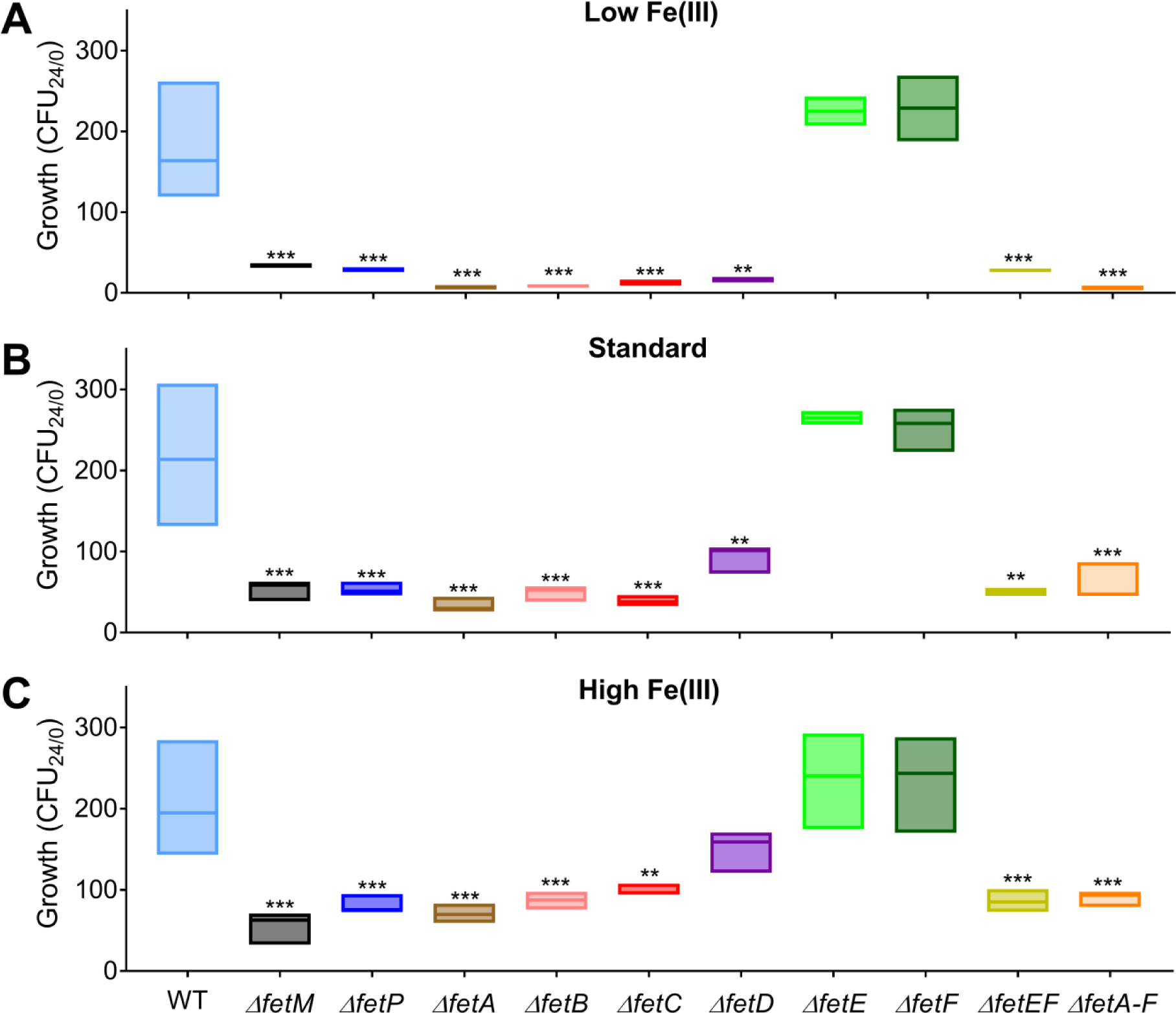
Growth monitored by CFU for *C. jejuni* gene deletion strains under different levels of iron availability. *C. jejuni* strains were cultured under low (MH + 5 µM DFO, **A**), standard (MH, **B**), or high (MH + 100 µM FeCl_3_, **C**) Fe(III) availability, with growth monitored by CFU. Corresponding results for *C. jejuni* complemented strains are in **Supplemental Figure S3**. Each strain was assayed in triplicate except for wild-type (WT), which was cultured for every growth experiment and hence was assayed with 18 biological replicates. CFU/mL was determined for each culture by dilution plating (5 technical replicates). CFU/mL values calculated for each culture at 24 h were divided by the CFU/mL at 0 h to represent the amount of growth in each culture (CFU_24/0_). Box plots for show the median and range. Statistical comparison for deletion mutants versus wild-type was performed using the Student’s *t*-test: * = *p*<0.05, ** = *p*<0.01, *** = *p*<0.001.

Growth defects demonstrated by gene deletion mutants were fully restored to that of wild-type by ectopic chromosomal complementation. Overall, the trends in growth observed by OD_600_ (**Figure 2 and Supplemental Figure S2**) were consistent with those observed by CFU (**Figure 3 and Supplemental Figure S3**), with complementation diminishing the possibility that these phenotypes resulted from polar effects on downstream genes. As a further measure of validation, the phenotypes observed for the *C. jejuni* strains that were being used as experimental standards (Δ*fetABCDEF*, *fetABCDEF*^C^, Δ*fetP*, and *fetP*^C^) were generally consistent with those in the original studies (10, 11).

Gene deletion strains Δ*fetM*, Δ*fetP*, Δ*fetA*, Δ*fetB*, Δ*fetC*, Δ*fetD*, and Δ*fetABCDEF* exhibited significant growth defects compared to wild-type by OD_600_ at 24 h and 30 h under all levels of iron availability. Additionally, growth after 30 h for Δ*fetP*, Δ*fetA*, Δ*fetB*, Δ*fetC*, Δ*fetD*, and Δ*fetABCDEF* strongly correlated with the level of iron availability for each strain (40-65% reductions with 5 µM DFO, 32-205% increases with 100 µM FeCl_3_). Growth of Δ*fetM* varied little upon iron restriction or supplementation (7% reduction with 5 µM DFO, 17% increase with 100 µM FeCl_3_), indicating the highest degree of insensitivity to iron availability in the absence of *fetM* gene compared to the other gene deletions (**Supplemental Figure S4**). These trends for growth defects and sensitivity to iron availability for Δ*fetM*, Δ*fetP*, Δ*fetA*, Δ*fetB*, Δ*fetC*, Δ*fetD*, and Δ*fetABCDEF* were consistent with CFU data, with the exception that high iron availability was sufficient to restore the growth of Δ*fetD* to a level that was similar to wild-type (*p* = 0.06).

By both OD_600_ and CFU, individual Δ*fetE* and Δ*fetF* strains did not demonstrate growth defects and were similarly insensitive to iron availability when compared to wild-type. The double deletion mutant Δ*fetEF*, however, had significantly reduced growth by OD_600_ and CFU/mL compared to wild-type, Δ*fetE*, and Δ*fetF* under all levels of iron availability. Growth at 30 h for Δ*fetEF* was dependent iron on availability (46% reduction with 5 µM DFO, 49% increase with 100 µM FeCl_3_).

### Streptomycin sensitivity of *C. jejuni* strains

The biphasic phenotype exhibited by wild-type *C. jejuni* under increasing concentrations of the antibiotic streptomycin is lost upon deletion of the *fetABCDEF* gene cluster (**Supplemental Figure S5**) (10). To investigate the role of each gene in biphasic streptomycin resistance, all deletion strains were assayed for minimum inhibitory concentration (MIC) of streptomycin (**Supplemental Figure S5**). Control wild-type *C. jejuni* cultures exhibited streptomycin-sensitive growth from 0 to 1 µg/mL streptomycin and streptomycin-tolerant growth from 1 to 4 µg/mL streptomycin.

All *C. jejuni* gene deletion strains demonstrated increased sensitivity to streptomycin. Δ*fetM*, Δ*fetA*, Δ*fetB*, Δ*fetC*, Δ*fetD,* Δ*fetABCDEF,* and Δ*fetP* exhibited unimodal growth and hence a loss of biphasic phenotype (**Supplemental Figure S5A**). At 1 µg/mL streptomycin, Δ*fetM*, Δ*fetA*, and Δ*fetD* demonstrated minimal growth similar to Δ*fetABCDEF* (7–10% of no-streptomycin control), Δ*fetB* and Δ*fetP* demonstrated an intermediate phenotype (23% of no-streptomycin control), and Δ*fetC* was similar to wild-type (33% of no-streptomycin control). At 2 and 4 µg/mL streptomycin, Δ*fetM*, Δ*fetP*, Δ*fetA*, Δ*fetB*, Δ*fetC*, and Δ*fetD* all showed similar growth to Δ*fetABCDEF* (2–6% of no-streptomycin control) (**Supplemental Figure S5A**), whereas Δ*fetE*, Δ*fetF*, and Δ*fetEF* demonstrated intermediate growth, between that of wild-type and Δ*fetABCDEF* (**Supplemental Figure S5B**). At 8 and 16 µg/mL streptomycin, all strains showed minimal growth. Wild-type biphasic phenotype and MIC were restored in all complemented strains (**Supplemental Figures S5C and S5D**).

### Expression of *C. jejuni* FetA protein is iron-regulated and independent of FetBCDEF

Due to the strong iron-dependent growth defect observed upon *fetA* deletion and the high level of *fetA* conservation in homologues of the *fet* gene cluster, *fetA* was selected as a gene to characterize further. To examine the protein expression levels of FetA under standard vs. iron-limited conditions, a 2xFlag-tagged version of FetA was expressed in the Δ*fetA* and Δ*fetABCDEF* deletion strains using the pRRC-based *fetA* complementation vector (pRRC_1651; **Supplemental Figure S1K**) under the control of the chloramphenicol resistance cassette promoter. FetA^2xFlag^ was able to restore the growth of Δ*fetA*, indicating that the tag had not disrupted function (**Supplemental Figure S6A**). FetA^2xFlag^ was unable to restore the growth of Δ*fetABCDEF*, which expectedly mimicked the growth defect phenotypes of the individual *fetB* to *fetD* deletion and *fetEF* double deletion strains. These strains were then analysed by western blot, probing for FetA with a monoclonal anti-Flag tag antibody (Coomassie-stained SDS-PAGE loading control: **Supplemental Figure S6B**; western blot: **Supplemental Figure S6C**). No FetA band was observed in the Δ*fetA* and Δ*fetABCDEF* controls. A band was observed for FetA in all tag-complemented strains, with higher protein levels under iron limitation. Full-length FetA is predicted to be ∼54 kDa but ran slightly smaller than expected by SDS-PAGE and produced a smeared band when visualized by western blotting, likely due to the large transmembrane region of this protein.

### *C. jejuni* FetE has capacity as a disulfide reductase

In light of our discovery that *fetE* and *fetF* function redundantly in iron scavenging, along with the prediction that these genes encode thioredoxins, FetE was selected for further characterization by functional assays and structural analysis. *C. jejuni* FetE was recombinantly expressed in *E. coli* BL21(DE3) and purified. Far-Western blot analysis (12) was used to screen for interactions between FetE, FetM, and FetP. While the expected interactions between FetM and FetP were observed, consistent with our previous work (12), no interactions of either FetM or FetP with FetE were detected (data not shown).

To verify whether *C. jejuni* FetE was capable of reducing disulfide bonds, an insulin disulfide reduction assay was selected as a standard method of thioredoxin characterization (26). The alpha and beta chains of insulin are linked by two disulfide bonds that can be reduced to precipitate the free beta chain. This produces an increase in absorbance at 650 nm that correlates to the rate of disulfide reduction, where baseline insulin reduction by dithiothreitol (DTT) can be increased by the addition of proteins with disulfide reductase activity. This assay was performed for *C. jejuni* FetE with comparison to a standard thioredoxin, *E. coli* Trx, and a blank sample (no protein added) representing baseline insulin reduction (**Figure 4**).

**Figure 4:**
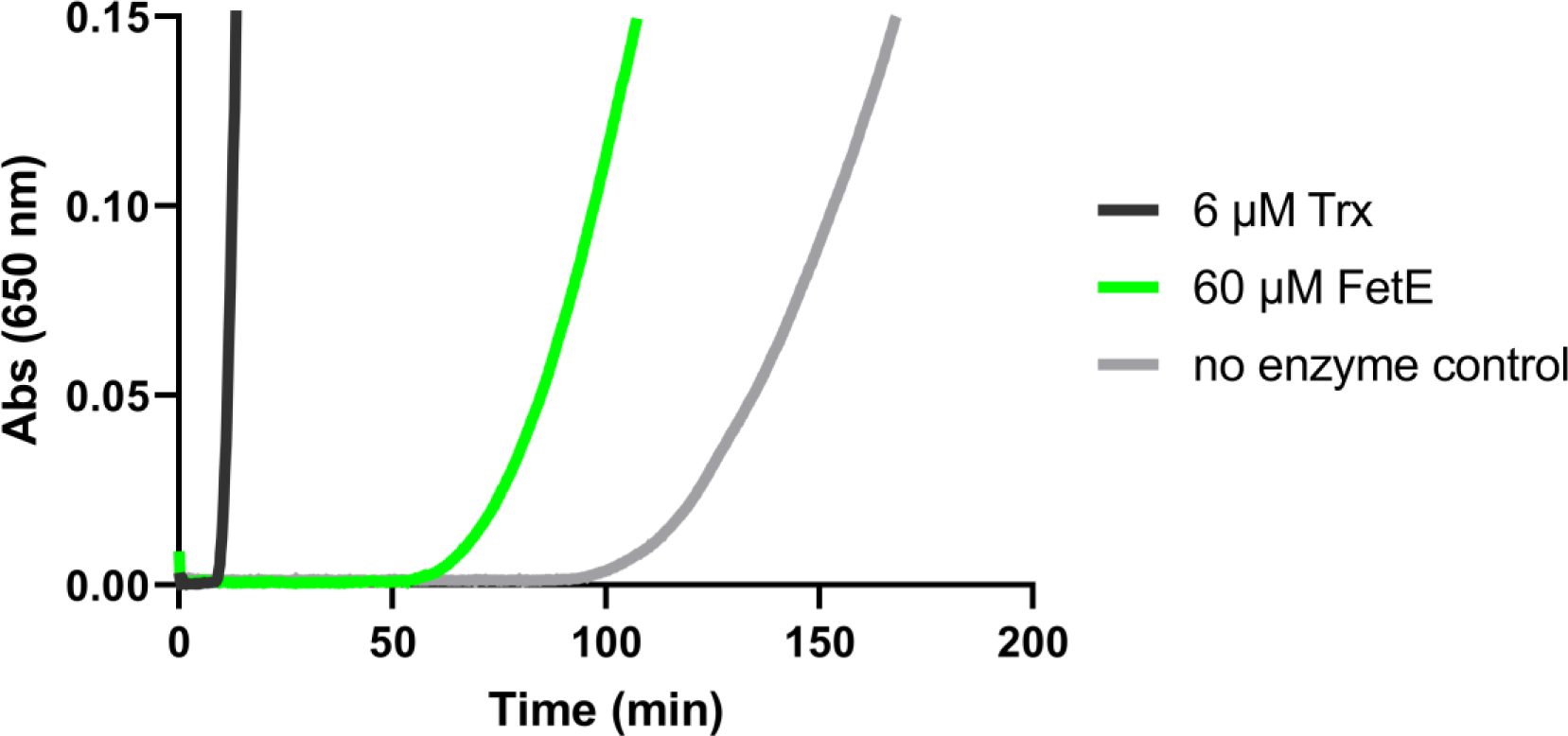
*C. jejuni* FetE exhibits disulfide reductase activity. Reduction of an intermolecular disulfide bond in bovine insulin (0.13 mM prepared in 0.1 M potassium phosphate pH 7.0, 2 mM EDTA, and 0.33 mM DTT) was monitored by an increase in absorbance at 650 nm.

Addition of *E. coli* Trx (6 uM) drastically increased the rate of insulin reduction well above the disulfide reductase activity of baseline (no protein), with the rate observed here for *E. coli* Trx being similar to those observed in previous studies (26). Addition of *C. jejuni* FetE (60 uM) also increased disulfide reduction above baseline, albeit to a lesser extent than *E. coli* Trx. Hence, these results indicate a capacity of *C. jejuni* FetE to mediate disulfide reduction.

### *C. jejuni* FetE is structurally related to thioredoxins

The crystal structure of a soluble construct of FetE lacking the lipobox was solved as a monomer to 1.50 Å resolution. FetE consisted of a 5-stranded β-sheet with 3 α-helices on one side and one short α-helix on the other. A structural similarity search of FetE against representative protein folds (PDB25) using the DALI server (27) revealed highest similarity to characterized proteins such as *Streptococcus gordonii* thiol-disulfide oxidoreductase SdbA (the top hit with Z-score 17.6; r.m.s.d. 2.1 Å over 133 Cα atoms; PDB ID: 5UM7) and *Mycobacterium tuberculosis* DsbE (Z-score 15.5; r.m.s.d. 2.2 Å over 127 Cα atoms; PDB ID: 1LU4). SdbA is an oxidase involved in the formation of disulfide-bonded proteins (28). Similarly, Mtb DsbE functions as an oxidase, which is atypical compared to the reductase role of its Gram-negative DsbE counterparts (29). Other top DALI hits remain uncharacterized. Alignment of 30 unique FetE homolog sequences (E-value cutoff = 0.0001) mapped onto the surface of the FetE crystal structure using Consurf (30) revealed conservation of the predicted key catalytic CXXC motif, which are the most highly conserved residues on the surface of FetE (**Supplemental Figure S7**).

### Deletion of *fetEF* affects DTNB reduction by *C. jejuni* cell-free extracts

A colorimetric 5,5’-dithiobis-(2-nitrobenzoic acid) (DTNB) reduction assay was used to monitor for differences in disulfide reduction capacity between extracts of *C. jejuni* wild-type, Δ*fetE*, Δ*fetF*, and Δ*fetEF*. DTNB consists of two aromatic groups linked by a disulfide bond, with disulfide reduction resulting in production of a thiol anion that absorbs strongly at 412 nm. Cell-free extracts were prepared for *C. jejuni* wild-type, Δ*fetE*, Δ*fetF*, and Δ*fetEF*, normalised for total protein concentration, then incubated with DTNB and NADH in a cuvette for spectrophotometric determination of the disulfide reduction rate (**Figure 5**).

**Figure 5:**
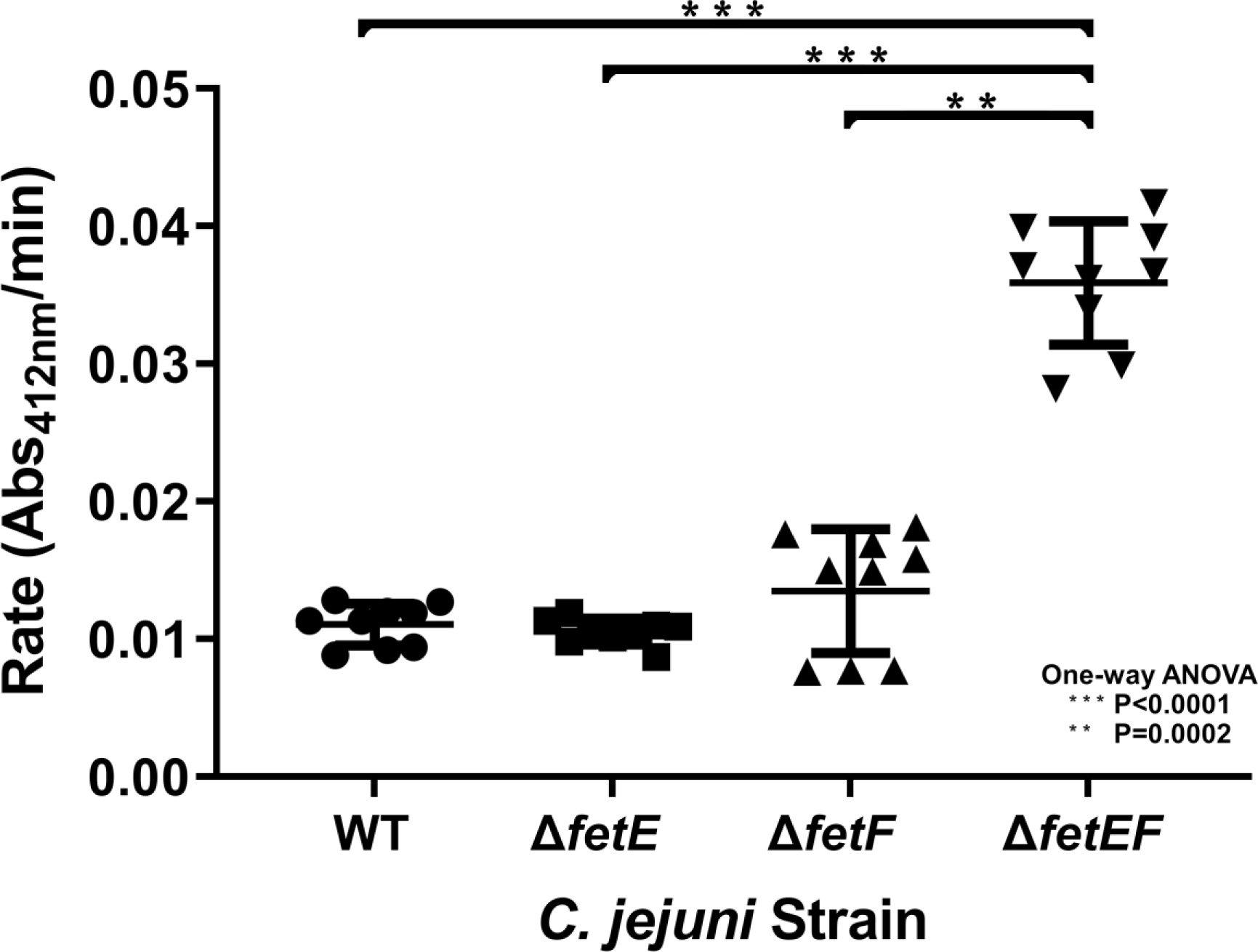
DTNB reduction by *C. jejuni* wild-type, Δ*fetE*, Δ*fetF*, and Δ*fetEF* cell extracts. The overall rate for each strain represents extracts from three mid-log phase cultures incubated for 3 h under iron limitation (5 µM DFO). Reduction of 0.1 mM DTNB in 0.2 mM NADPH, 50 mM Tris-HCl pH 7.2 by each extract was performed three times, with error bars representing standard deviation and statistical significance determined by one-way ANOVA.

Similar DTNB reduction rates were observed for cell extracts from *C. jejuni* wild-type (0.0110 ± 0.0014 Abs_412_/min), Δ*fetE* (0.0105 ± 0.0009 Abs_412_/min), and Δ*fetF* (0.0135 ± 0.0042 Abs_412_/min) strains. Comparatively, *C. jejuni* Δ*fetEF* cell extract showed a significantly higher rate of DTNB reduction (0.0359 ± 0.0042 Abs_412_/min).

## DISCUSSION

In addressing the extensive global morbidity and socioeconomic burden caused by *C. jejuni*, it is of high priority to gain a deeper understanding of the molecular systems that support virulence during infection, such as the upregulated *fetMP-fetABCDEF* gene cluster. In this study, we systematically deconstructed the *fetMP-fetABCDEF* gene cluster to assess the role of each gene in *C. jejuni* growth during iron scarcity as well as the functionality of one of the predicted oxidoreductase proteins.

Strong growth defects and high sensitivity to iron availability were observed for Δ*fetA*, Δ*fetB,* Δ*fetC,* and Δ*fetD*. These phenotypes mirrored a deletion of the cluster (Δ*fetABCDEF*), indicating that each individual gene product was important for growth in general, with defects exacerbated under iron restriction and partially rescued by the addition of exogenous iron to the medium. Moreover, these strains showed sensitivity to iron availability comparable to that of Δ*fetP*, which plays a key role in iron transport (12), highlighting an equally important role for the individual FetA and FetBCD proteins in iron uptake. This is reinforced by the high conservation of equivalent genes encoding predicted membrane permeases and ABC transporters in all known homologues of the *fetMP-fetABCDEF* cluster across several bacterial phyla (10).

The observation of higher FetA protein levels under iron limitation across all 2xFlag-tag-complemented strains indicates that FetA expression is iron-dependent. The FetA^2xFlag^ construct included the intergenic region between *fetP* and *fetA,* with *fetA* expression under the control of a constitutively-expressing Cm promoter. As the Cm promoter it not known to be iron regulated, this suggests the presence of regulatory elements either in the intergenic region between *fetP* and *fetA* or within *fetA* itself. Alternatively, *fetA* may be post-transcriptionally regulated. Additionally, the presence of detectable FetA in Δ*fetABCDEF^2xFlag-fetA^* indicates that expression and proper folding of FetA is independent of the downstream *fet* genes.

ABC transporters are a common component of bacterial iron uptake systems, with ATP hydrolysis often driving the passage of a Fe-siderophore complex from the periplasm to the cytoplasm through a channel formed by the two transmembrane proteins (31). If FetMP-FetABCDEF represents one iron uptake system in which FetM is the sole iron permease, then the function (and substrate) for the putative ABC transporter encoded by *fetBCD* remains unclear. Despite homology between *fetB* and *fetC* (28% sequence identity), deletion of either gene resulted in a strong growth defect, indicating that both genes are required for transporter function. This suggests the specific requirement of a FetB-FetC heterodimer for proper function of this gene cluster, and that FetB-FetB or FetC-FetC homodimers are either not formed or cannot sufficiently restore the growth defects of Δ*fetB* or Δ*fetC*.

Individual deletion *fetE* or *fetF* in *C. jejuni* did not correspond to a growth defect under any level of iron availability, whereas the deletion of both genes (Δ*fetEF*) resulted in a growth defect in iron-restricted medium. This suggests that *fetE* and *fetF* perform redundant functions to support *C. jejuni* growth during iron restriction, as the presence of either gene is sufficient to maintain growth comparable to wild-type. Interestingly, while increased sensitivity to iron availability generally coincided with greater sensitivity to streptomycin, suggesting that sufficient iron is important in resisting antibiotic effects, the streptomycin sensitivity of Δ*fetE* and Δ*fetF* was equivalent to that of Δ*fetEF,* and all three deletion strains were intermediate compared to the other deletion strains. *fetE* and *fetF* are homologs (22% sequence identity) predicted to encode periplasmic, membrane-associated protein disulfide reductases. We demonstrated that FetE contains a thioredoxin fold and can reduce insulin disulfide. However, the comparatively slow rate of insulin reduction by FetE compared to *E. coli* Trx, the structural similarity to the thiol-disulfide oxidases SdbA and DsbE, and the higher rate of DTNB reduction by extracts of *C. jejuni* Δ*fetEF* than those of wild-type, Δ*fetE*, and Δ*fetF,* suggests that the primary role of FetE and FetF may be to act as oxidases *in vivo*. This would be consistent with our previous observation that *C. jejuni* Δ*fetABCDEF* had greater survival than wild-type upon exposure to oxidative stress (10), which suggested that the presence of FetABCDEF increased susceptibility to the deleterious effects of oxidation.

Other than the individual thioredoxin deletion strains Δ*fetE* and Δ*fetF*, every tested deletion strain exhibited growth defects. While no strain was completely devoid of growth, the deletion of *fetM* resulted in a growth defect which was the least correlated with iron availability. All other deletion strains with growth defects exhibited greater growth restoration upon increased iron availability. For the Fet cluster, only homologs of FetM have to date been shown to directly transport iron (17, 18). This suggests that other iron uptake systems in *C. jejuni* are responsible for the partial growth observed upon deletion of any one non-thioredoxin *fet* gene. However, these same uptake systems are only able to minimally improve growth of the Δ*fetM* strain under iron supplementation, supporting its key role in direct iron transport. The other *fet* genes, with our proposed role in preparing iron for transport, would exhibit a growth effect that is more dependent on iron availability.

Overall, the experimental results described here have advanced understanding on the collective roles of the Fet system components in relation to *C. jejuni* iron scavenging. In combining these new findings with an *in silico* investigation and prior literature, we propose an updated model for how the system encoded by *fetMP-fetABCDEF* may function (**Figure 6**). In this revised model, an iron-chelator complex first passes through the outer membrane via an as yet unidentified transporter. Upon entering the periplasm, the iron is released from the chelator and transported into the cytoplasm by the cooperative action of the iron binding protein FetP and the iron permease FetM. Our previous studies in *C. jejuni* and uropathogenic *E. coli* strongly suggest an iron oxidation/reduction-based mechanism for iron transport (11, 12, 18). Based on our studies presented here, we predict that FetE and FetF play overlapping roles in supporting FetMP-based iron transport by actively relaying the necessary reducing power likely sourced from FetB, FetC, and FetD, functioning as a single heteromeric ABC transporter, and through FetA. In the absence of FetM, a major route for iron to cross the inner membrane, growth is impaired irrespective of the presence of other Fet proteins, making this strain more resistant to growth recovery upon iron supplementation. FetA, FetBCD and FetEF, conversely, each critically support the redox dependency of FetMP iron uptake function. Hence deletion of components from FetABCDEF result in the growth of FetMP-intact strains that is more dependent on overall iron availability. Despite being a double-edged sword, as FetABCDEF increases susceptibility to oxidative stress (10), we demonstrate that this cluster is conserved because it plays an important role in cell growth in conjunction with FetMP.

**Figure 6:**
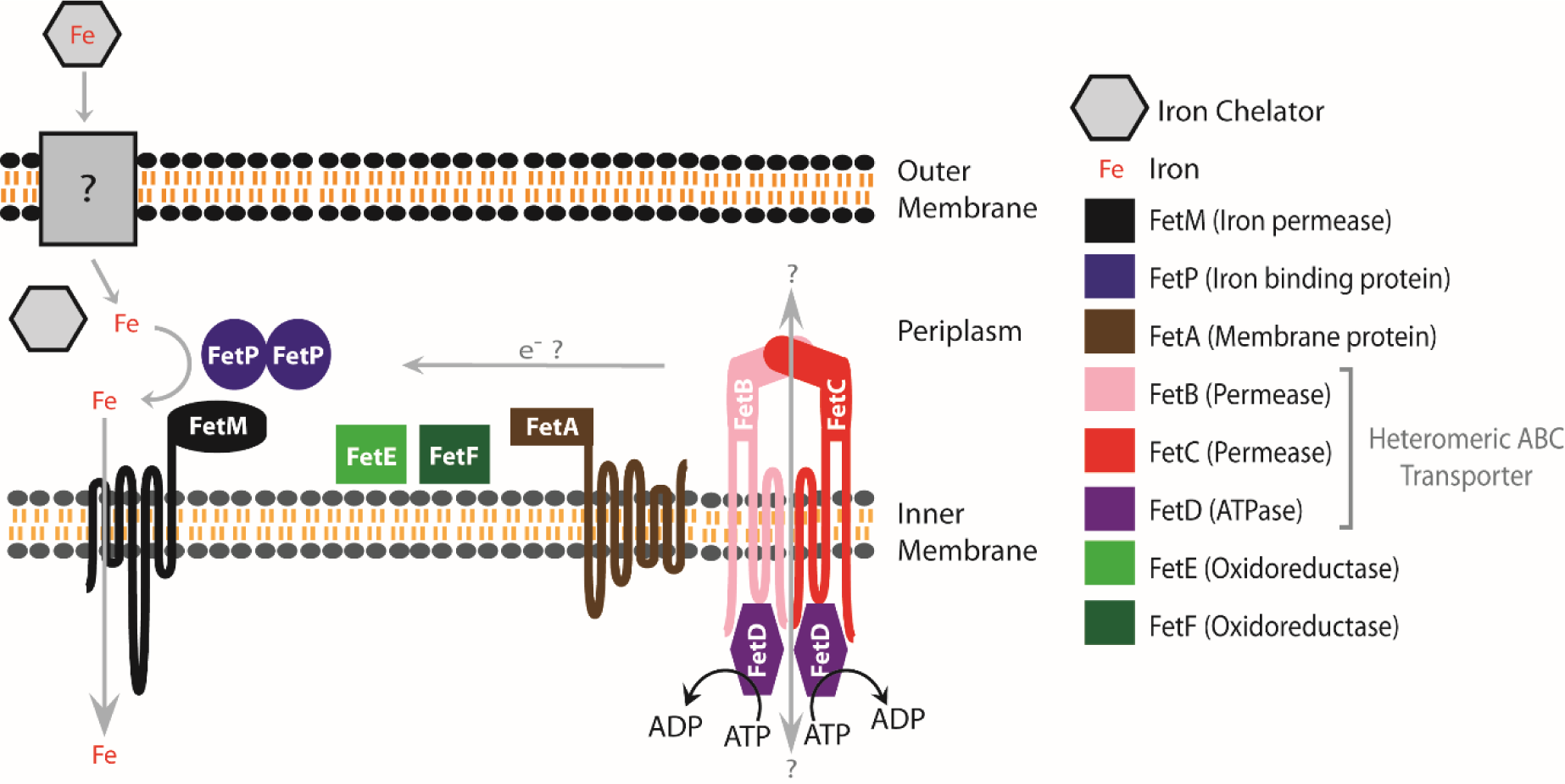
Model of the FetMP-FetABCDEF system based on predicted and known functions in iron transport. Adapted and updated from Liu, et al (10).

## CONCLUSION

The significant global disease burden caused by *C. jejuni* has provided impetus for research into novel systems important for pathogenesis and pathogenesis-related attributes. The *fetMP-fetABCDEF* genes have stood out as highly upregulated during human infection and in the presence of human fecal extracts, with only recent work bringing the importance of the downstream cluster *fetABCDEF* to light. This study has addressed gaps in knowledge relating to the *C. jejuni fetMP-fetABCDEF* gene cluster through a combined microbial genetics, molecular biology, biochemistry, and structural biology approach. All components of this gene cluster emerged as determinants of growth during iron scarcity, a known virulence-determining factor during *C. jejuni* infection. Expression of the integral membrane protein FetA was shown to be iron-regulated and independent of FetBCDEF. FetBCD likely forms a heteromeric ABC transporter essential to the function of the Fet cluster. *C. jejuni* FetE most closely resembles the structure of thiol-disulfide oxidases and demonstrated comparatively poorer disulfide reduction activity. Additionally, cell extracts from the deletion of *fetEF* exhibited the greatest reduction activity, together suggesting that FetE and FetF may function as oxidases *in vivo*.

## METHODS

### Design and construction of *C. jejuni* gene deletion and complemented strains

For gene deletion strains, the wild-type gene along with flanking regions was cloned into a pGEM-T plasmid and a portion of the gene was replaced with a non-polar *aphA3* kanamycin (Km) resistance cassette. Natural transformation of the modified pGEM suicide vector into *C. jejuni* allowed replacement of the target gene by homologous recombination at the flanking regions (**Supplemental Figure S1A-I**). For complemented strains, the wild-type gene was cloned into pRRC, which was naturally transformed into its respective *C. jejuni* mutant strain for integration of the gene and an upstream chloramphenicol (Cm) resistance cassette at one or more of three ectopic loci in the chromosome (32) (**Supplemental Figure S1J,K**).

A list of all strains, plasmids and primers used during construction for each strain are provided in **Supplemental Table S1**. *C. jejuni* 81-176 (clinical isolate from diarrheic patient) was used as the wild-type strain for all experiments (33). Plasmids and strains were verified by PCR analysis followed by Sanger sequencing (Genewiz). Growth conditions for *C. jejuni* and *E. coli*, detailed strain construction protocols, and determination of total iron content in the standard medium by ICP-MS are described in the **Supplemental Methods**.

### *C. jejuni* growth experiments for sensitivity to iron availability

All *C. jejuni* strains were grown overnight on MH-TV agar plates with Km (deletion strains) or Cm (complemented strains), streaked onto fresh equivalent plates, then grown for another 6 h. Cells were harvested and resuspended in MH-TV broth (10 mL) to an OD_600_ of 0.0004 (WT), 0.002 (all complemented strains), 0.005 (Δ*fetE*, Δ*fetF*), or 0.02 (all other deletion mutants) to consistently achieve cells in mid-log-phase (OD_600_ of 0.3–0.6) after a further 18 h of shaking incubation (200 rpm). Mid-log-phase cultures were resuspended in fresh 2×MH-TV and then dispensed into 96 well plates containing equivalent volume aqueous solutions of DFO (10 μM), water, or FeCl_3_ (200 μM) to achieve 200 μL 1×MH-TV starting cultures at an initial OD_600_ of 0.005, corresponding to low iron (MH-TV + 5 μM DFO), standard (MH-TV), and high iron (MH-TV + 100 μM FeCl_3_) conditions. Throughout incubation, growth was monitored by OD_600_ (Thermo Fisher Scientific Varioskan Flash plate reader) at 0 h, 6h, 24 h, and 30 h, and by CFUs at 0 h and 24 h. CFU/mL values calculated for each culture at 24 h were divided by the CFU/mL at 0 h to represent the amount of growth in each culture (CFU_24/0_). All strains were assessed with three biological replicates for each level of iron availability, and CFUs were determined using five technical replicates. Statistical differences were calculated using the Student’s *t*-test.

### *C. jejuni* growth experiments for sensitivity to streptomycin

Full experimental details are described in the **Supplemental Methods**.

### Expression of 2xFlag-tagged FetA in *C. jejuni*

A C-terminal 2x repeat Flag-tag was inserted into *C. jejuni fetA* by FastCloning (34) using the constitutively-expressing complementation vector pRRC_1651 as template (**Supplemental Figure S1K**) and Q5 DNA polymerase. pRRC_1651 includes 93 C-terminal bp of *fetP,* 82 bp of the intergenic region and the complete 1404 bp *fetA* gene. *C. jejuni* strains Δ*fetA and* Δ*fetABCDEF* were then complemented with this construct following the method to generate Δ*fetA^c^*, producing Δ*fetA^2xFlag-FetA^* and Δ*fetABCDEF^2xFlag-FetA^.* Successful insertion was confirmed by sequencing of purified genomic DNA. Methods to confirm proper functionality of the tagged FetA variant in the deletion strains are described in the **Supplemental Methods**.

To examine the expression of tagged FetA in *C. jejuni*, the variant-complemented strains were grown to mid-log-phase in 15 ml MH-TV, resuspended in fresh MH-TV supplemented with or without 10 μM DFO, and pelleted after 3 h of growth. The harvested cell pellets were analysed by SDS-PAGE and probed by western blot with an anti-Flag antibody (Genscript A01868).

### Expression of *C. jejuni* FetE and other Fet proteins

The expression vector for *C. jejuni* FetE was synthesized by GeneArt (Invitrogen) into pET151/D-TOPO. The DNA sequence encoded an N-terminal 6xHis tag, a V5 epitope, a TEV cut site, and residues IDPFT followed by amino acids 22–162 of native FetE (CJJ81176_1655). This vector was transformed into *E. coli* BL21(DE3) for expression and purification, following the protocol for *E. coli* FetA with slight modifications. Cells were induced with 0.25 mM IPTG, lysed in 30 mM Tris, 200 mM NaCl, 5 mM imidazole and 2 mM TCEP, pH 7.5 and dialysed into 30 mM Tris, 100 mM NaCl, and 2 mM TCEP, pH 7.5 after His-tag removal with TEV protease. TEV protease was removed with a second nickel resin purification step using dialysis buffer and concentrated. *C. jejuni* FetM and FetP were prepared as part of other studies (11, 12).

### FetE crystallization and structure determination

FetE was crystallized in space group P2_1_ under 1.8 M ammonium sulfate and 0.1 M sodium acetate. Crystals were then soaked in 0.06 M sodium acetate, 1.6 M ammonium sulfate, and 0.5 M sodium iodide, cryoprotected using 30% glycerol, flash frozen with liquid nitrogen and sent to the SSRL beamline 9-2. A 1.95 Å resolution dataset was collected at 1.6 Å wavelength and iodide phasing was used to successfully solve the crystal structure. This structure was then used for molecular replacement against a non-iodide soaked 1.5 Å resolution dataset collected at CLS beamline CMCF-ID. Data collection and refinement statistics are summarized in **Supplemental Table S2**. The coordinates and observed structure factor amplitudes have been deposited in the PDB under the accession code 8T4C.

### *C. jejuni* cell extracts and DTNB reduction assay

Protocols for the colorimetric DTNB reduction assay with *C. jejuni* cell extracts were adapted from methods used by Kaakoush *et al.* (2007) to identify *C. jejuni* protein disulfide reductases (35). *C. jejuni* cell extract preparation and the adapted assay are described in the **Supplemental Methods**.

## ACKNOWLEDGEMENTS

The authors acknowledge the significant contributions and mentorship of the late Professor Erin Gaynor to this work and we are grateful for Dr. Gaynor’s family’s support in seeing this work published. We also thank Martha Liu for providing constructive input to the project and Jina Yeom for her preliminary work on the streptomycin sensitivity assays. We thank Jenny Wallace and Emilisa Frirdich for training with designing deletion and complementation strains, as well as training for *C. jejuni* growth assays. This research was supported by grants from the Michael Smith Foundation for Health Research (MSFHR) to T.R-S. (RT 18437), and Canadian Institutes of Health Research (CIHR) to E.C.G. (MOP-68981) and an Natural Science and Engineering (NSERC) Discovery Grant to MEPM (RGPIN-2022-04568). Use of the Stanford Synchrotron Radiation Lightsource, SLAC National Accelerator Laboratory, is supported by the U.S. Department of Energy, Office of Science, Office of Basic Energy Sciences under Contract No. DE-AC02-76SF00515. The SSRL Structural Molecular Biology Program is supported by the DOE Office of Biological and Environmental Research, and by the National Institutes of Health, National Institute of General Medical Sciences (P30GM133894). The contents of this publication are solely the responsibility of the authors and do not necessarily represent the official views of NIGMS or NIH. Part of the research described in this paper was also performed using beamline CMCF-ID at the Canadian Light Source, a national research facility of the University of Saskatchewan, which is supported by the Canada Foundation for Innovation (CFI), the Natural Sciences and Engineering Research Council (NSERC), the National Research Council (NRC), the Canadian Institutes of Health Research (CIHR), the Government of Saskatchewan, and the University of Saskatchewan. The authors declare no conflicts of interest.

## Supplemental Methods

### 1. *C. jejuni* growth conditions

Unless otherwise indicated, *C. jejuni* was grown at 38 °C under microaerobic and capnophilic conditions (6% O_2_, 12% CO_2_) produced by a tri-gas incubator (Sanyo) or CampyGen system (Oxoid). The growth medium was MH (Oxoid lot number 2429934) broth or agar plates (1.7% w/v agar) with vancomycin (10 μg/mL) and trimethoprim (5 μg/mL) (MH-TV), and kanamycin (Km, 50 μg/mL) or chloramphenicol (Cm, 20 μg/mL) as necessary.

### 2. *E. coli* growth conditions

*E. coli* cultures used in plasmid construction were grown at 37 °C under atmospheric conditions using Luria-Bertani broth or agar plates (1.5% w/v agar). *E. coli* growth media were supplemented with Km (25 μg/mL), Cm (20 μg/mL), ampicillin (Ap, 100 μg/mL), or 5-bromo-4-chloro-3-indolyl-β-D-galactopyranoside (X-gal, 80 μg/mL), where appropriate.

### 3. *C. jejuni* strain construction

Construction of gene deletion strains Δ*fetM*, Δ*fetA*, Δ*fetB*, Δ*fetC*, Δ*fetD*, Δ*fetE*, and Δ*fetF* was achieved by replacing a portion of each gene with the non-polar *aphA3* Km resistance (Km^R^) cassette from pUC18-K2 (1). Each gene with approximately 300-400 bp flanking regions was amplified by PCR from *C. jejuni* 81-176 genomic DNA. A polyA tail was added to the PCR product before ligation to a pGEM-T vector (Promega) and transformation into *E. coli* DH5α. Constructs harbouring the resulting plasmid (pGEM_*gene*) were selected using Ap and X-gal plates. pGEM_*gene* was then used as a template for inverse PCR using primers with XbaI and KpnI restriction sites engineered into the 5’ ends. The inverse PCR product was digested with XbaI and KpnI and ligated to the *aphA3* Km^R^ cassette similarly digested out of pUC18-K2 to give pGEM_Δ*gene*. After transformation into *E. coli* DH5α, constructs containing pGEM_Δ*gene* were selected using Km plates. pGEM_Δ*gene* acts as a suicide vector in *C. jejuni*, incorporating into the chromosome by homologous recombination to replace the target gene *via* the identical 300–400 bp flanking regions. *C. jejuni* 81-176 was naturally transformed with pGEM_Δ*gene* as previously described (2), and Km^R^ colonies with the gene of interest disrupted by *aphA3* were selected.

Construction of complemented strains *fetM^C^*, *fetA^C^*, *fetB^C^*, *fetC^C^*, *fetD^C^*, *fetE^C^*, *fetF^C^*, and *fetEF^C^* was achieved by complementing each deletion mutant with the relevant gene(s) at an ectopic locus in the chromosome. The relevant gene(s) were amplified by PCR from *C. jejuni* 81-176 genomic DNA using primers with XbaI and MfeI restriction sites engineered into the 5’ end. In cases where XbaI or MfeI sites could not be used, sites were engineered for the isoschizomers SpeI or EcoRI, respectively. The PCR product was ligated to a similarly digested pRRC vector (3) to give pRRC_*gene*, which was transformed into *E. coli* DH5α. Constructs containing pRRC_*gene* were selected using Cm plates. pRRC is an integration vector that inserts the cloned gene along with an upstream Cm resistance (Cm^R^) cassette into one of three 16S ribosomal regions in the *C. jejuni* genome (3). The pRRC_*gene* plasmid was naturally transformed into the corresponding *C. jejuni* gene deletion strain (Δ*gene*). Constructs with both the gene deletion (Km^R^) and complementation (Cm^R^) were selected on Km and Cm plates. *C. jejuni* strains Δ*fetEF* and *fetEF*^C^ were constructed by these same protocols using the upstream primers for *fetE* with the downstream primers for *fetF*.

### 4. Determining total iron content of standard medium by ICP-MS

ICP-MS was used to measure the amount of ^56^Fe in the standard medium (MH-TV) that was used in all *C. jejuni* growth experiments for this study. Three separate batches of standard medium were prepared and each subsampled (500 µL) for use as a replicate. Closed vessel sample digestion was performed in 35% HNO_3_ at 110 °C on a hotplate. Solvent was removed by drying before samples were redissolved in 1% HNO_3_ with ^45^Sc (20 ppb) as an internal standard. ICP-MS was performed using a NexION 300D (Perkin Elmer) equipped with a SC-2 DX autosampler, DXi-FAST micro-peristaltic pump, a cyclonic spray chamber, a triple cone interface, a quadrupole ion deflector and Universal Cell Technology. Calibration was performed using the IV-Stock-4 ICP calibration standard (Inorganic Ventures). All elements were run in reaction mode (using Dynamic Reaction Cell technology) using ammonia as a reaction gas to remove potential polyatomic interferences. The detection limit for ^56^Fe was determined as 0.356 ppb.

### 5. *C. jejuni* growth experiments for sensitivity to streptomycin

In assessing streptomycin sensitivity, trimethoprim and vancomycin were not added to MH at any stage. All *C. jejuni* strains were grown to mid-log-phase then resuspended in fresh 2×MH and dispensed into 96 well plates containing equivalent volumes of water or aqueous streptomycin solution at an initial OD_600_ of 0.02. This produced cultures in either unsupplemented medium (MH, positive control) or medium containing doubling concentrations of streptomycin (MH + 0.125–16 μg/mL streptomycin). After 48 h incubation, OD_600_ was determined using a Varioskan Flash plate reader (Thermo Fisher Scientific). Strains were assessed for all streptomycin concentrations with three biological replicates. Growth at each streptomycin concentration was expressed as a percentage of the positive control (unsupplemented MH) and graphed using GraphPad Prism 7.

### 6. Confirmation of proper functionality of 2xFlag-tagged FetA

Proper functionality of the tagged FetA variant in the deletion strains was determined by comparing growth of the variant-complemented strains to the strains complemented with native FetA and native FetABCDEF and their respective deletion strains, as in main Methods with modifications. Cells harvested from MH-TV plates were resuspended in MH-TV broth (3 mL) to an OD_600_ of 0.005 (Δ*fetA*, Δ*fetABCDEF* and Δ*fetABCDEF^2xFlag-fetA^*) or 0.0005 (Δ*fetA^c^*, Δ*fetABCDEF^c^* and Δ*fetA^2xFlag-fetA^*) for overnight starting cultures. Mid-log-phase cultures were resuspended in fresh MH-TV and then dispensed into 96 well plates with 0, 4, 8, 12, or 16 μM DFO to achieve 200 μL starting cultures at an initial OD_600_ of 0.0075. Growth was measured as OD_600_ (Thermo Fisher Scientific Varioskan Flash plate reader) after 24 h of incubation.

### 7. Insulin reduction assay

The insulin reduction assay protocol was adapted from established methods (4). A solution of potassium phosphate (0.1 M, pH 7.0) was prepared with EDTA (2 mM), insulin solution from bovine pancreas (0.13 mM, Sigma-Aldrich), and DTT (0.33 mM). The insulin reduction reaction was then started by addition of *C. jejuni* FetE (60 µM) or *E. coli* Trx (6 µM, Sigma-Aldrich). To measure baseline insulin reduction, phosphate buffer was added in place of the proteins.

### 8. Preparation of *C. jejuni* cell extracts and DTNB reduction assay

To prepare *C. jejuni* wild-type, Δ*fetE*, Δ*fetF*, and Δ*fetEF* strains for extract preparation, cultures (25 mL MH-TV) were grown as previously described to achieve robust mid-log-phase growth after 18 h of shaking incubation (200 rpm). At this stage, DFO (5 µM) was added to promote gene expression and cultures incubated for a further 3 h. Cells were pelleted (15000×g, 4 °C, 10 min), supernatant discarded, and each culture resuspended in 10 mL NaCl (150 mM) to wash the cells and increase cell density by a factor of 2.5. This wash procedure (pelleting then resuspension in 10 mL NaCl solution) was repeated three times before cell lysis by six cycles of freezing in liquid nitrogen and thawing. The cell-free extracts were then collected as the supernatant fraction after a final centrifugation step (15000×g, 4 °C, 10 min) and total protein concentration was determined by Bradford assay. To ensure reproducibility, three biological replicates (cultures) were used for each strain.

Prior to use in the DTNB reduction assay, each cell extract sample was normalized according to total protein concentration and divided into three technical replicates for the assay. 150 mM NaCl was used as the diluent and negative control. For the DTNB reduction assay, DTNB (0.1 mM) and NADPH (0.2 mM) were combined in 50 mM Tris-HCl (pH 7.2) in 1 cm path-length cuvettes. To commence DTNB reduction, normalized *C. jejuni* extracts were added to this mixture and change in absorbance at 412 nm was immediately recorded with a spectrophotometer over 1 min.

**Supplemental Figure S1:**
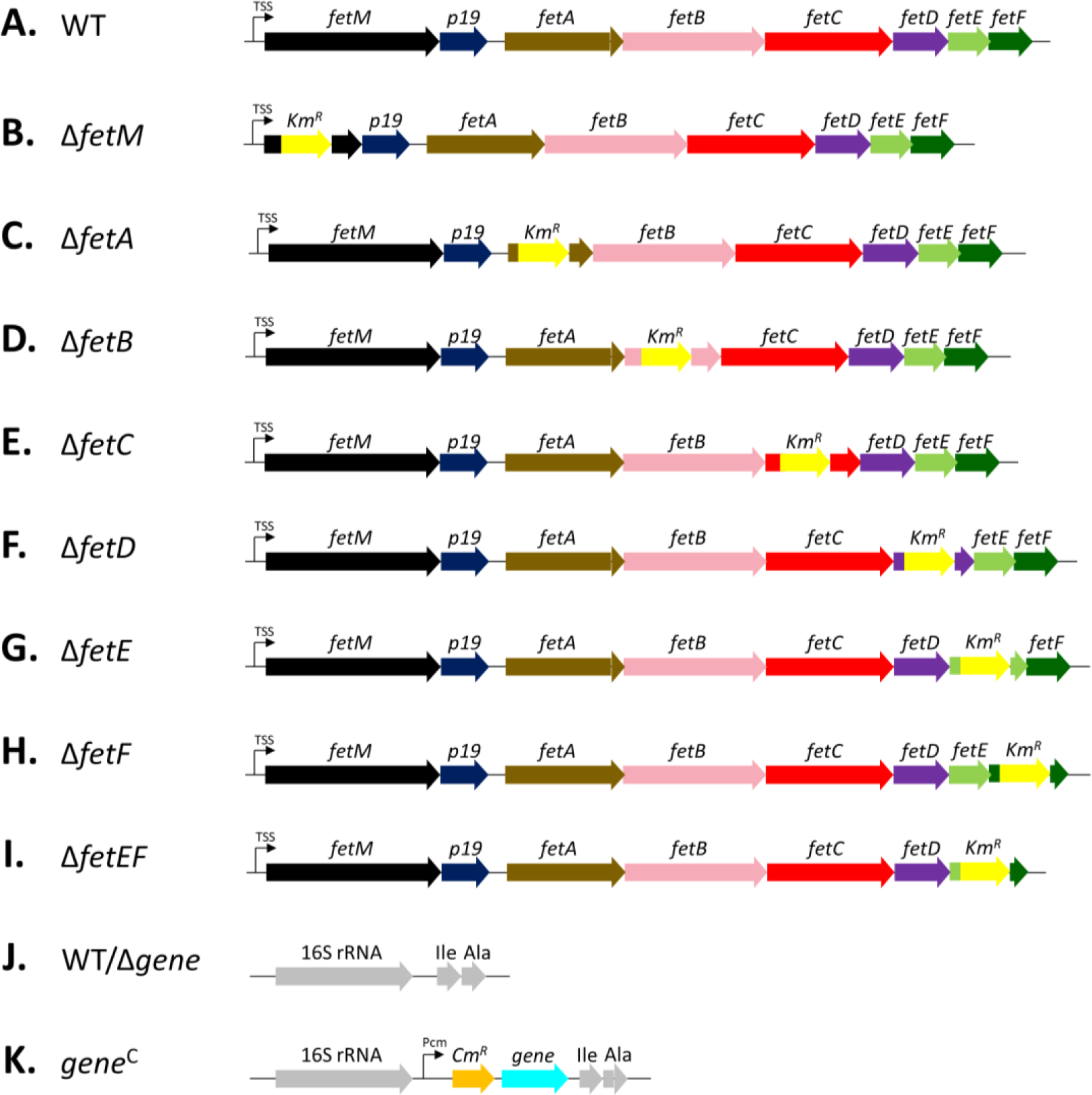
Construction of the *C. jejuni* gene deletion and complemented strains. (**A**) *C. jejuni* wild-type (WT) harbours the *1649*-*1656* gene cluster downstream of a transcription start site (TSS). (**B**-**I**) *C. jejuni* gene deletion mutants were constructed by replacing a portion of the target gene(s) with a kanamycin resistance cassette (Km^R^). (**J**) The 16S ribosomal RNA region of *C. jejuni* WT and uncomplemented deletion strains. Directly downstream of the gene encoding for 16S rRNA are genes for t-RNA^Ile^ (Ile) and t-RNA^Ala^ (Ala). (**K**) *C. jejuni* complemented strains were constructed by inserting a constitutively-expressing chloramphenicol promoter region (Pcm), chloramphenicol resistance cassette (Cm^R^), and the target gene into the 16S ribosomal RNA region of the corresponding deletion strain.

**Supplemental Figure S2:**
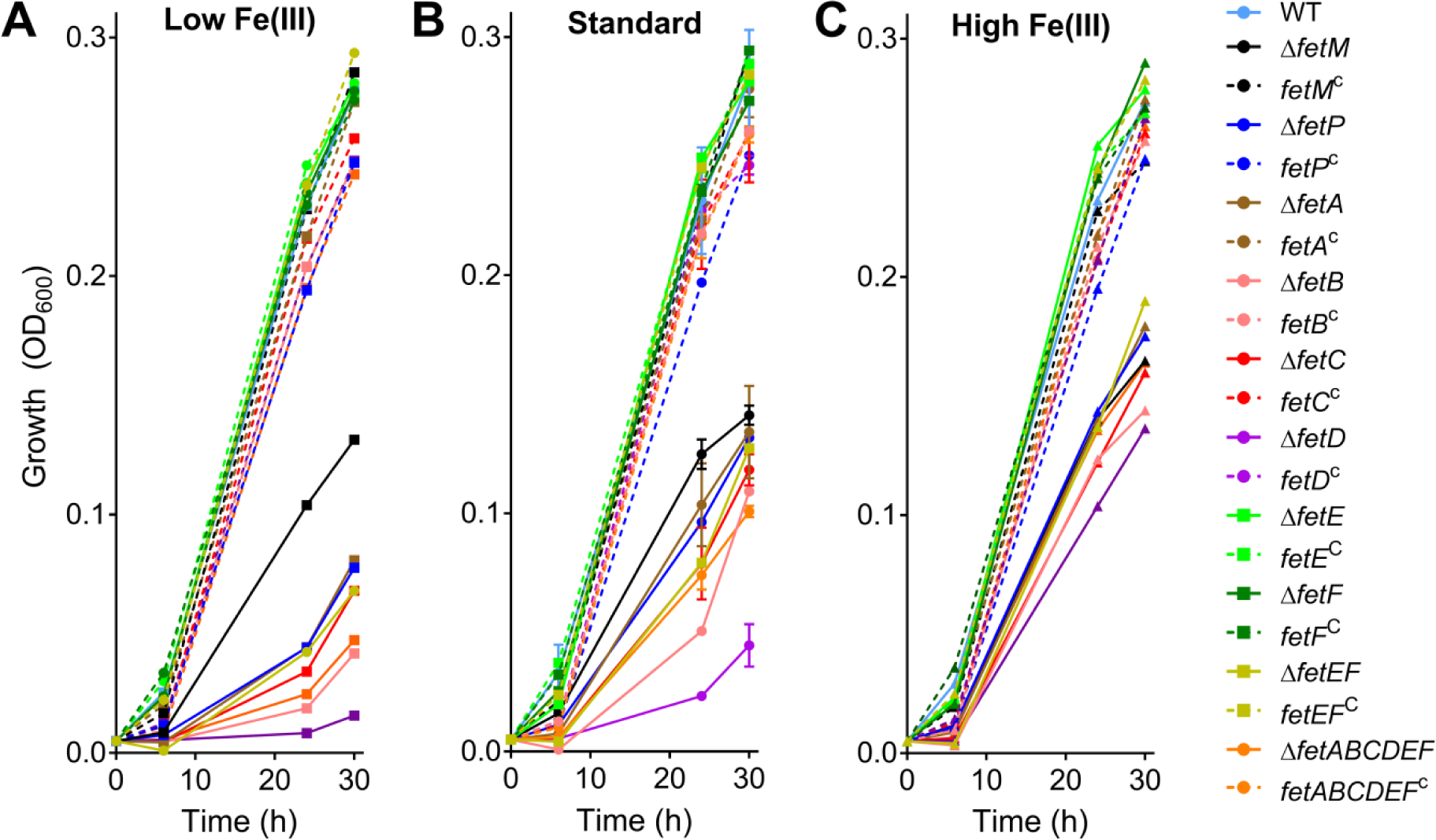
Growth by OD for *C. jejuni* gene deletion and complemented strains under different levels of iron availability. *C. jejuni* strains were cultured under low (MH + 5 µM DFO, **A**), standard (MH, **B**), or high (MH + 100 µM FeCl_3_, **C**) Fe(III) availability, with growth monitored by OD_600_. Each strain was assayed in triplicate except for wild type (WT), which was cultured for every growth experiment and hence was assayed with 18 biological replicates. Error bars represent standard deviation.

**Supplemental Figure S3:**
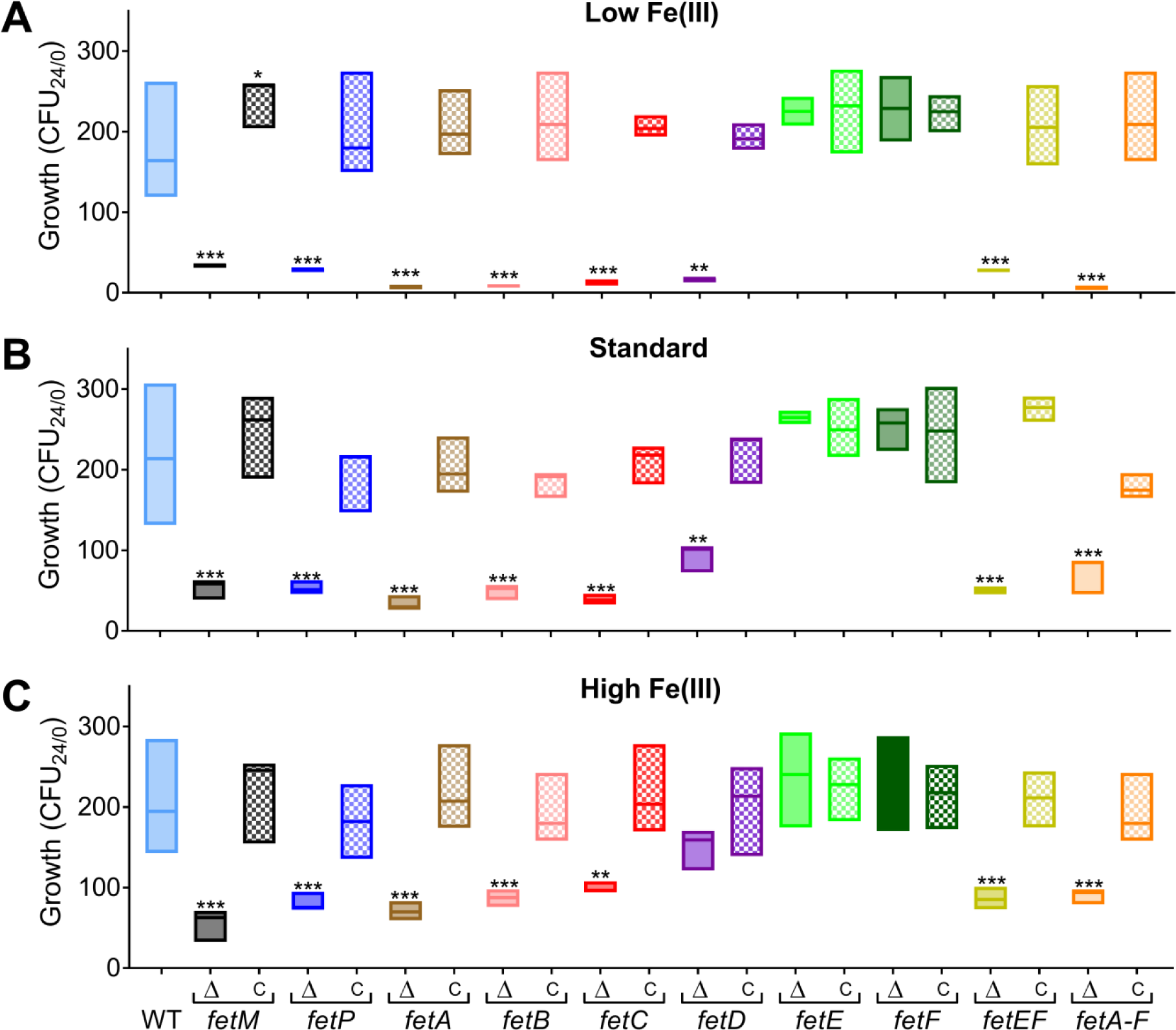
Growth by CFU for *C. jejuni* gene deletion and complemented strains under different levels of iron availability. *C. jejuni* strains were cultured under low (MH + 5 µM DFO, **D**), standard (MH, **E**), or high (MH + 100 µM FeCl_3_, **F**) Fe(III) availability, with growth monitored by CFU. Each strain was assayed in triplicate except for wild type (WT), which was cultured for every growth experiment and hence was assayed with 18 biological replicates. CFU/mL was determined for each culture by dilution plating (5 technical replicates). CFU/mL values calculated for each culture at 24 h were divided by the CFU/mL at 0 h to represent the amount of growth in each culture (CFU_24/0_). Box plots show the median and range. Statistical comparison for CFU_24/0_ of deletion mutants versus wild type was performed using the Student’s *t*-test: * = *p*<0.05, ** = *p*<0.01, *** = *p*<0.001.

**Supplemental Figure S4:**
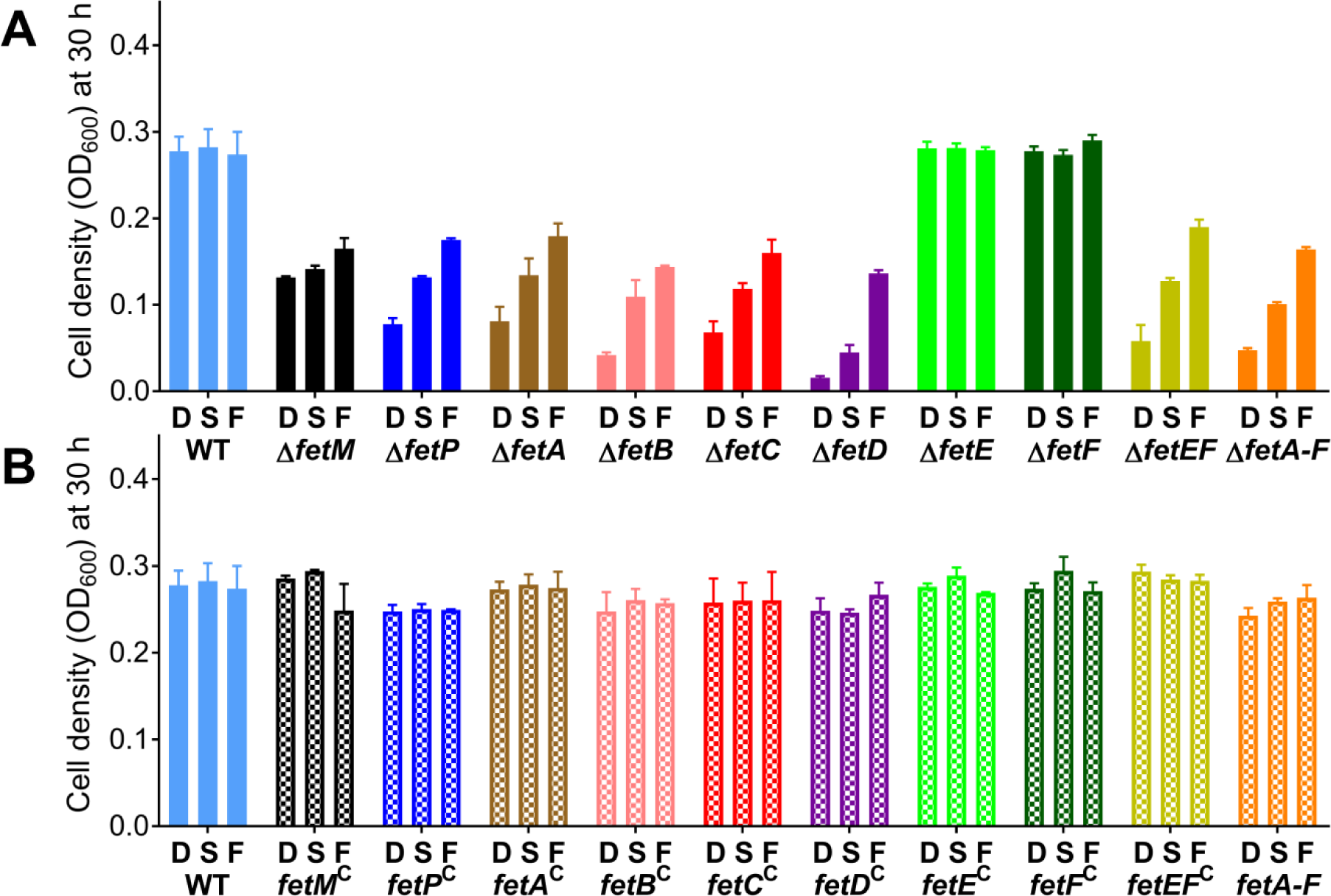
Iron sensitivity of *C. jejuni* gene deletion and complemented strains. This figure represents a subset of the data shown in Figure 2 and **Supplemental Figure S2**, showing cell density (by OD_600_) at 30 h of culturing *C. jejuni* wild type (WT), gene deletion (**A**), and complemented (**B**) strains under low (MH + 5 µM DFO, <u>D</u>), standard (MH, <u>S</u>), or high (MH + 100 µM FeCl_3_, <u>F</u>) Fe(III) availability. Each strain was assayed in triplicate except for wild type (WT), which was cultured for every growth experiment and hence was assayed with 18 biological replicates. Error bars represent standard deviation.

**Supplemental Figure S5:**
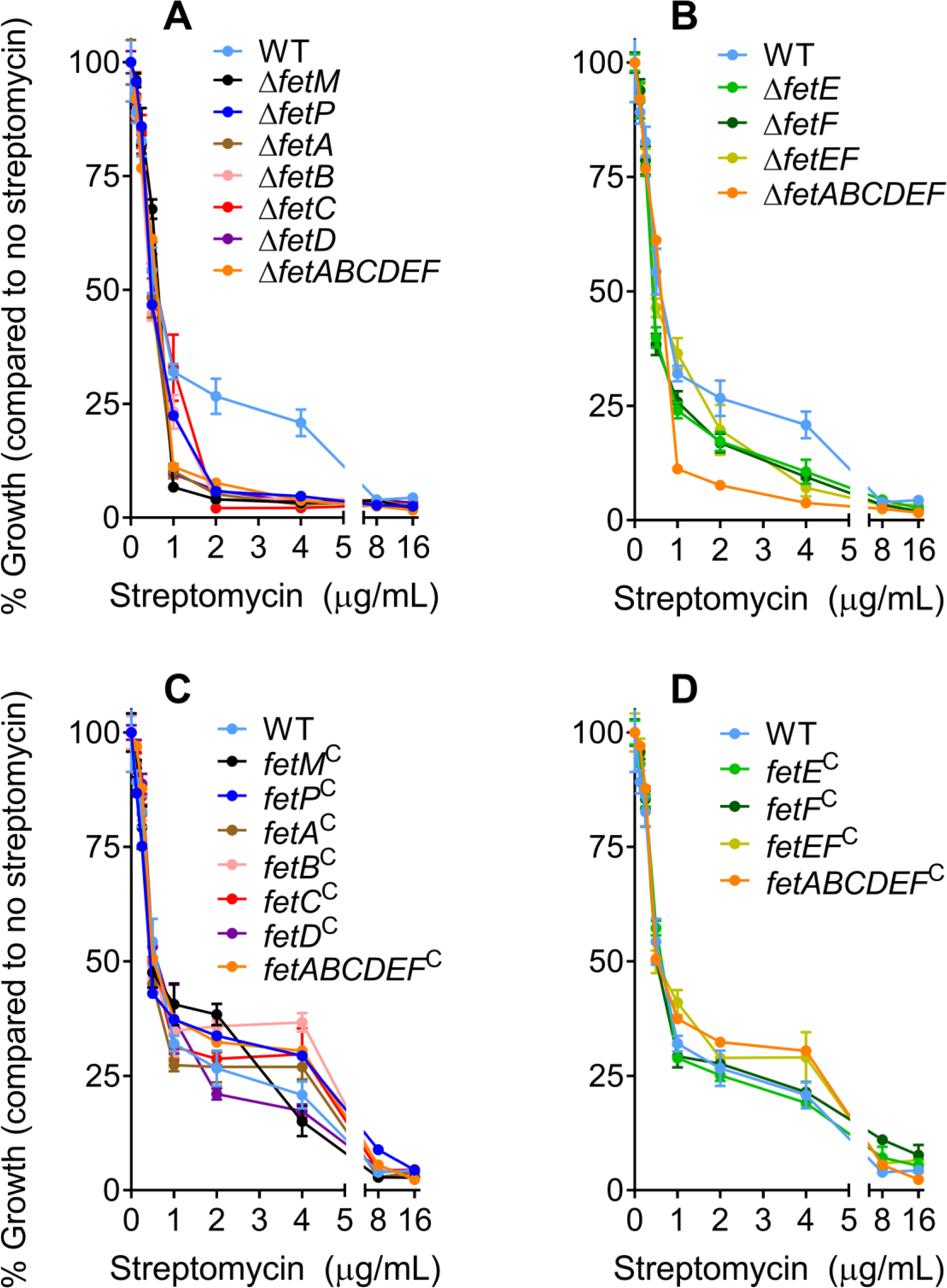
Streptomycin sensitivity for *C. jejuni* gene deletion and complementation strains. Deletion strains correspond to **A**) *fetM*, *fetP*, *fetA*, *fetB*, *fetC*, and *fetD*, and **B**) *fetE*, *fetF*, and *fetEF*, with comparison to wild-type (WT) and Δ*fetABCDEF*. Complementation strains correspond to **C**) *fetM*, *fetP*, *fetA*, *fetB*, *fetC*, and *fetD*, and **D**) *fetE*, *fetF*, and *fetEF*, with comparison to wild-type (WT) and Δ*fetABCDEF*. *C. jejuni* strains were cultured in standard medium (MH) with doubling concentrations of streptomycin. Growth was measured by OD_600_ at 48 h as a percentage of a no-streptomycin control. Error bars represent standard deviation from three different cultures.

**Supplemental Figure S6:**
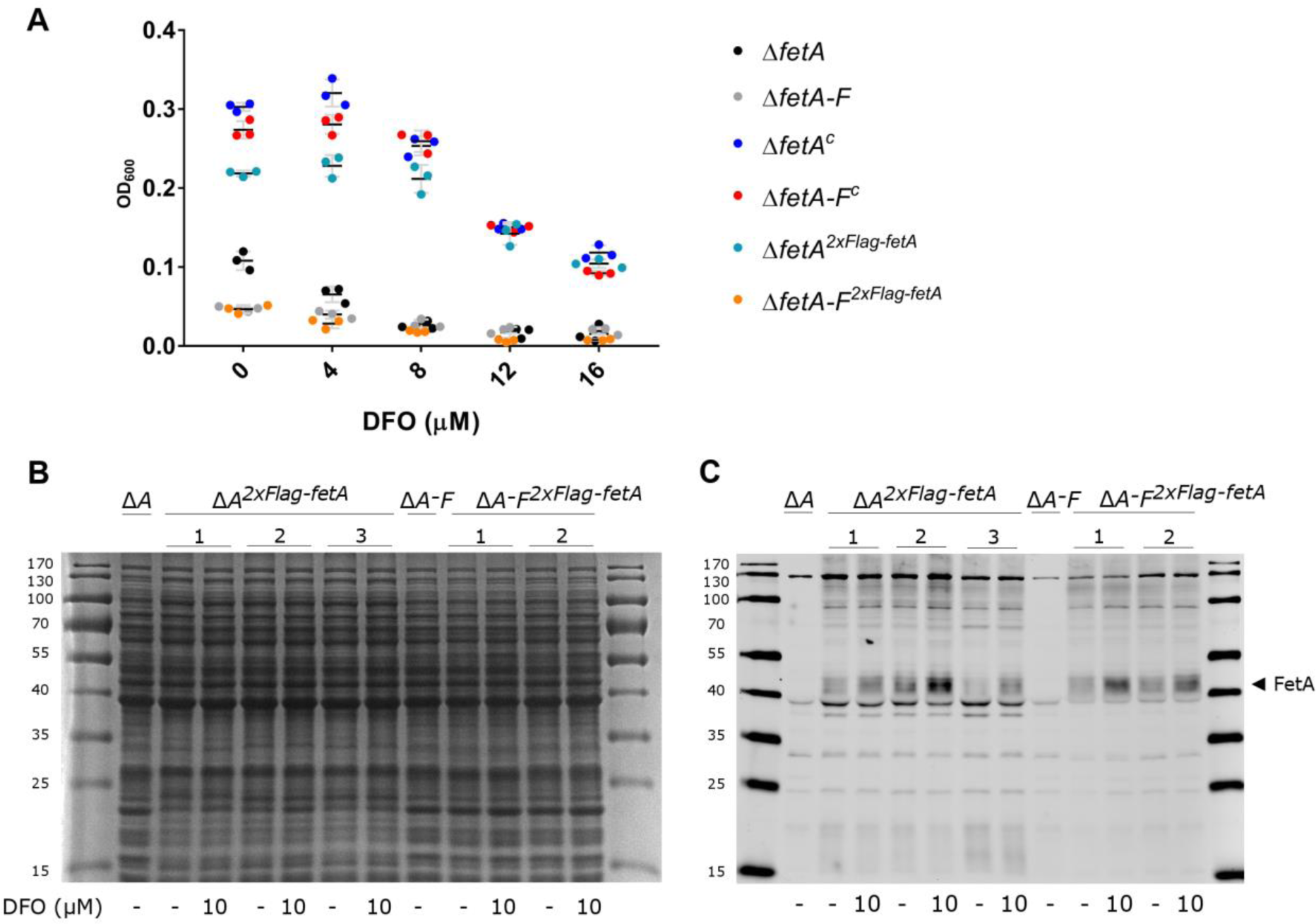
(A) The expression of 2xFlag-tagged FetA restores the growth of Δ*fetA* but not Δ*fetABCDEF*. Growth under different levels of iron availability were compared for strains containing a deletion in *fetA* (Δ*fetA*) or *fetABCDEF* (Δ*fetA-F*) and their complemented derivative strains (c = complemented with all deleted genes; 2xFlag-*fetA* = complemented with *fetA* containing a modified two-repeat C-terminal Flag-tag). Iron limitation was achieved by supplementation with increasing DFO. Each strain was assayed in triplicate (dots) with mean (black line) and standard deviation (grey line) shown. (B, C) The expression of FetA is iron-regulated. (B) is an SDS-PAGE gel to show equal sample loading amongst strains. (C) is a western blot using an anti-Flag antibody to demonstrate FetA levels in the deletion strains complemented with the Flag-tagged version of *fetA*. Cells were grown in MH-TV with either no iron limitation or with the addition 10 μM DFO (shown beneath the gels). Biological replicates for the complemented strains are numbered.

**Supplemental Figure S7:**
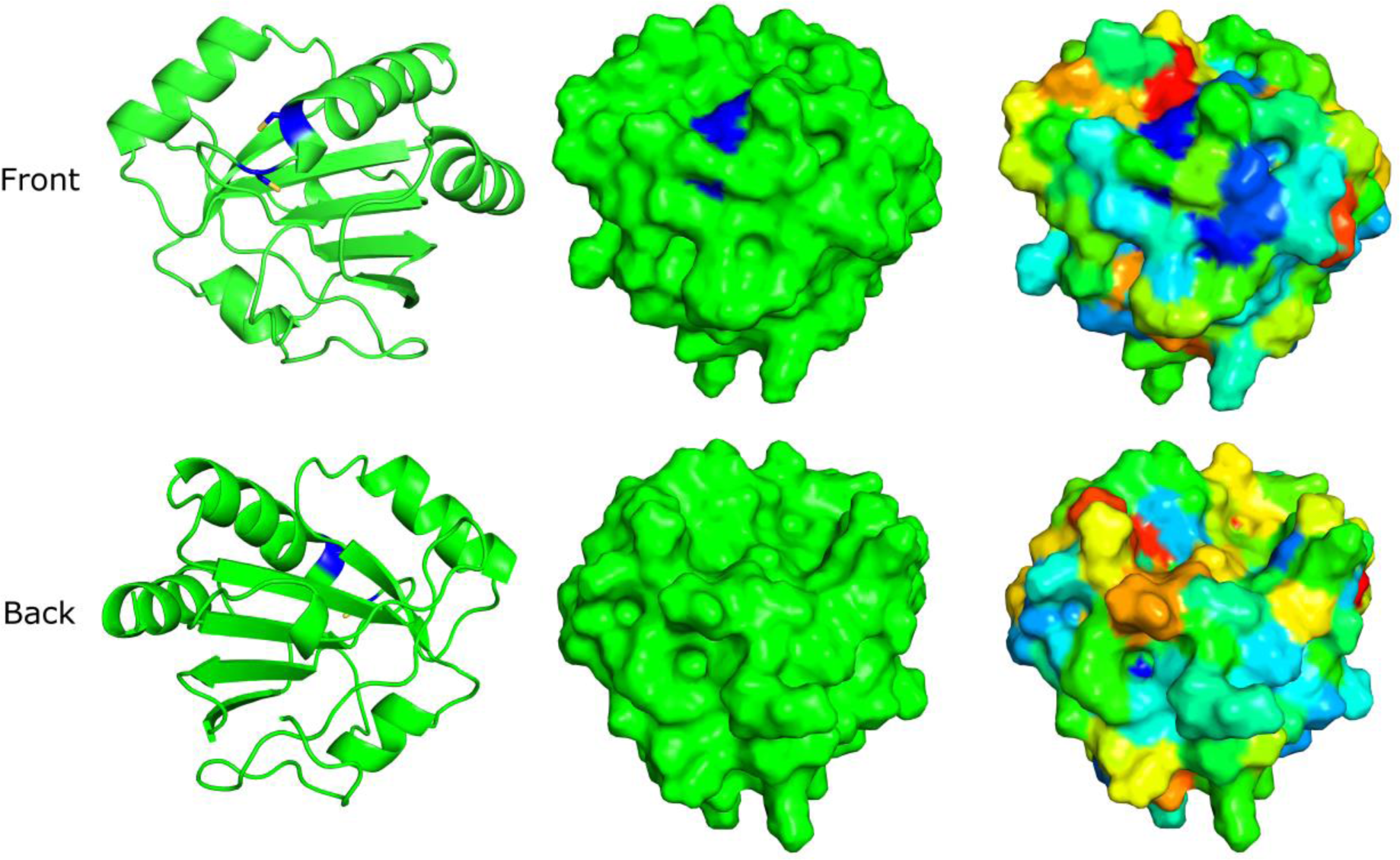
The crystal structure of FetE resembles thioredoxins. Cartoon (left) and surface (middle) representations of FetE with CXXC motif highlighted in blue to match conservation. Amino acid conservation analysis mapped to the structure of FetE (right), with most conserved residues in blue, followed by cyan, green, yellow, orange, and then least conserved residues in red.

**Supplemental Table S1:**
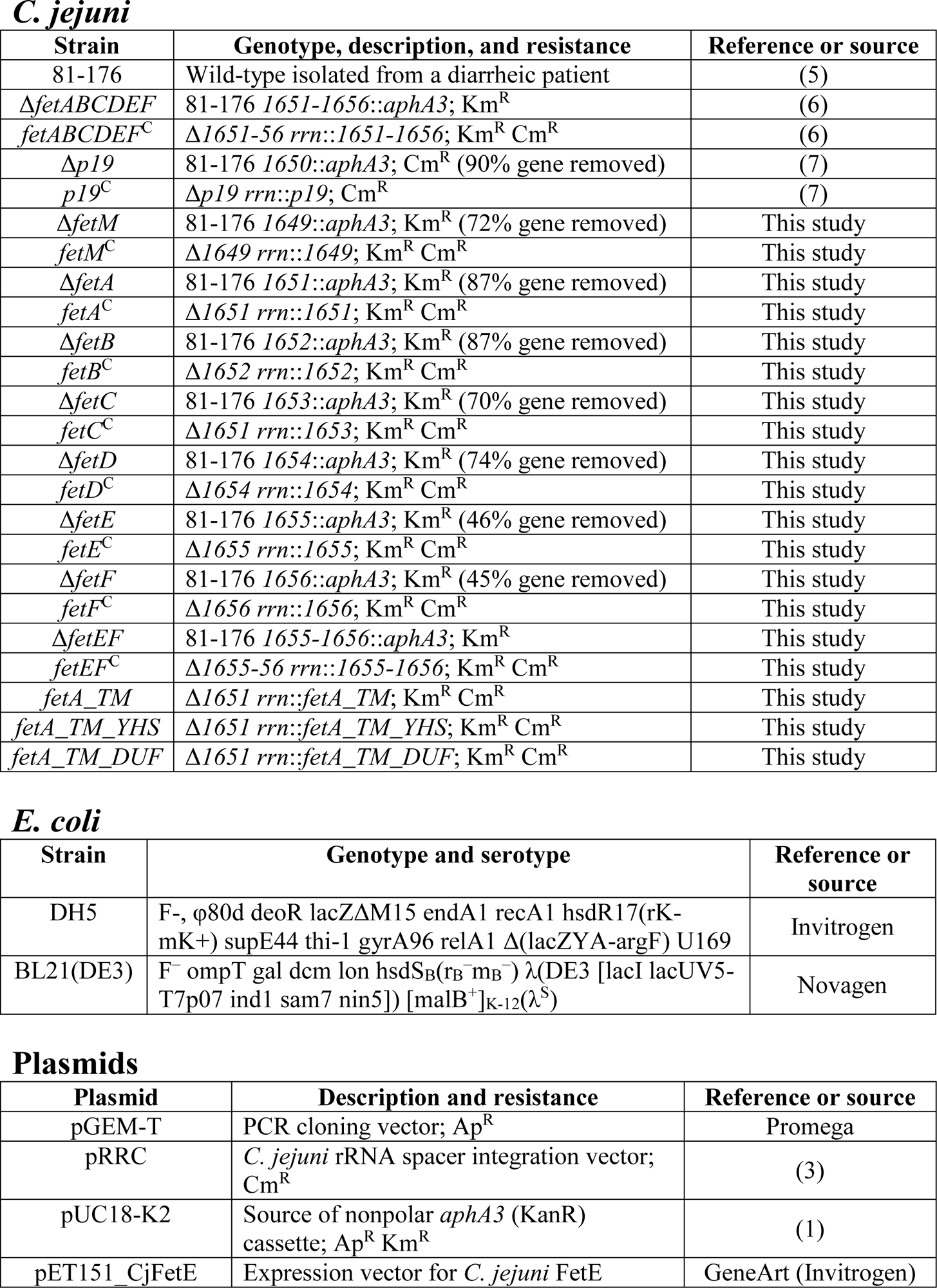
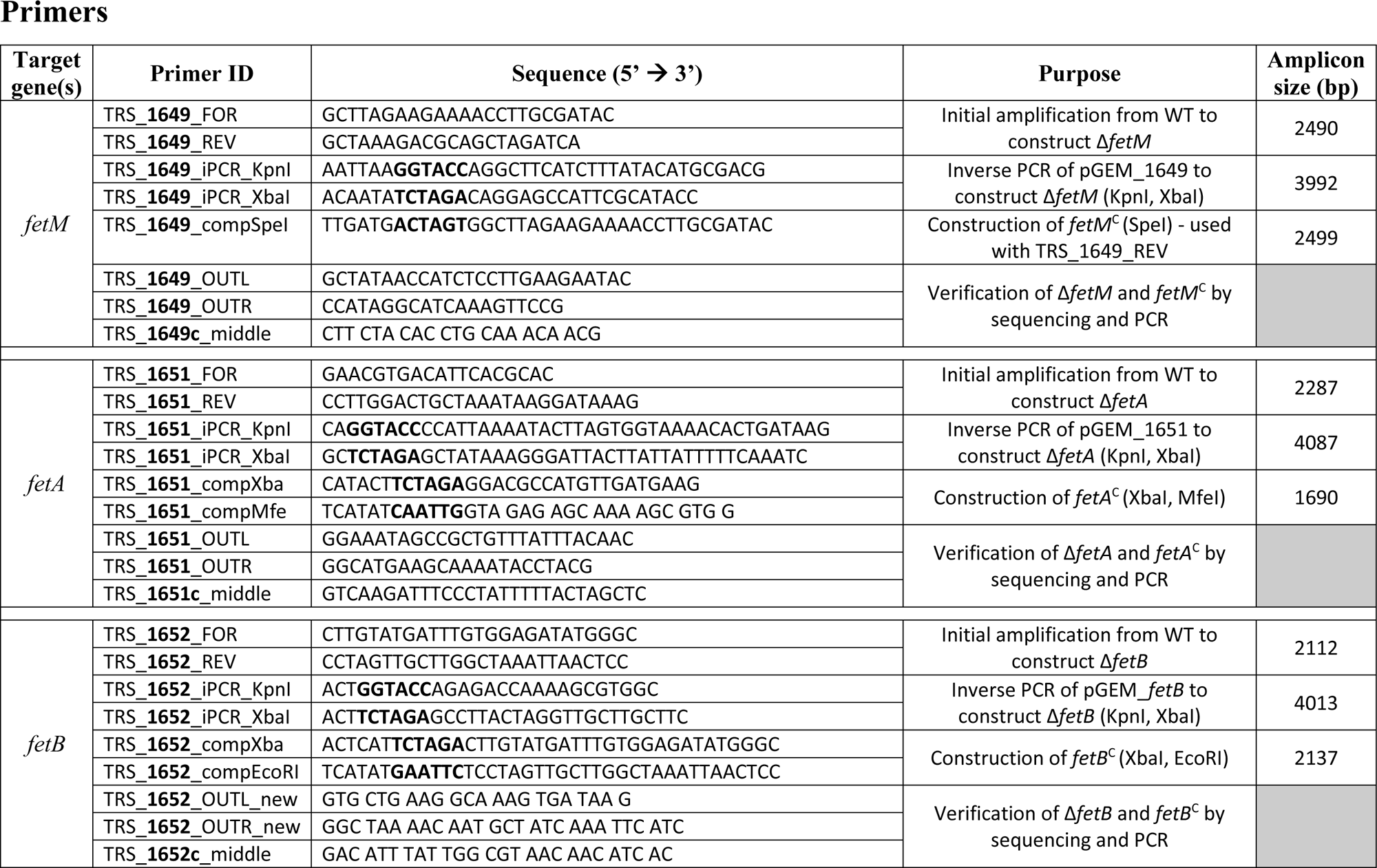
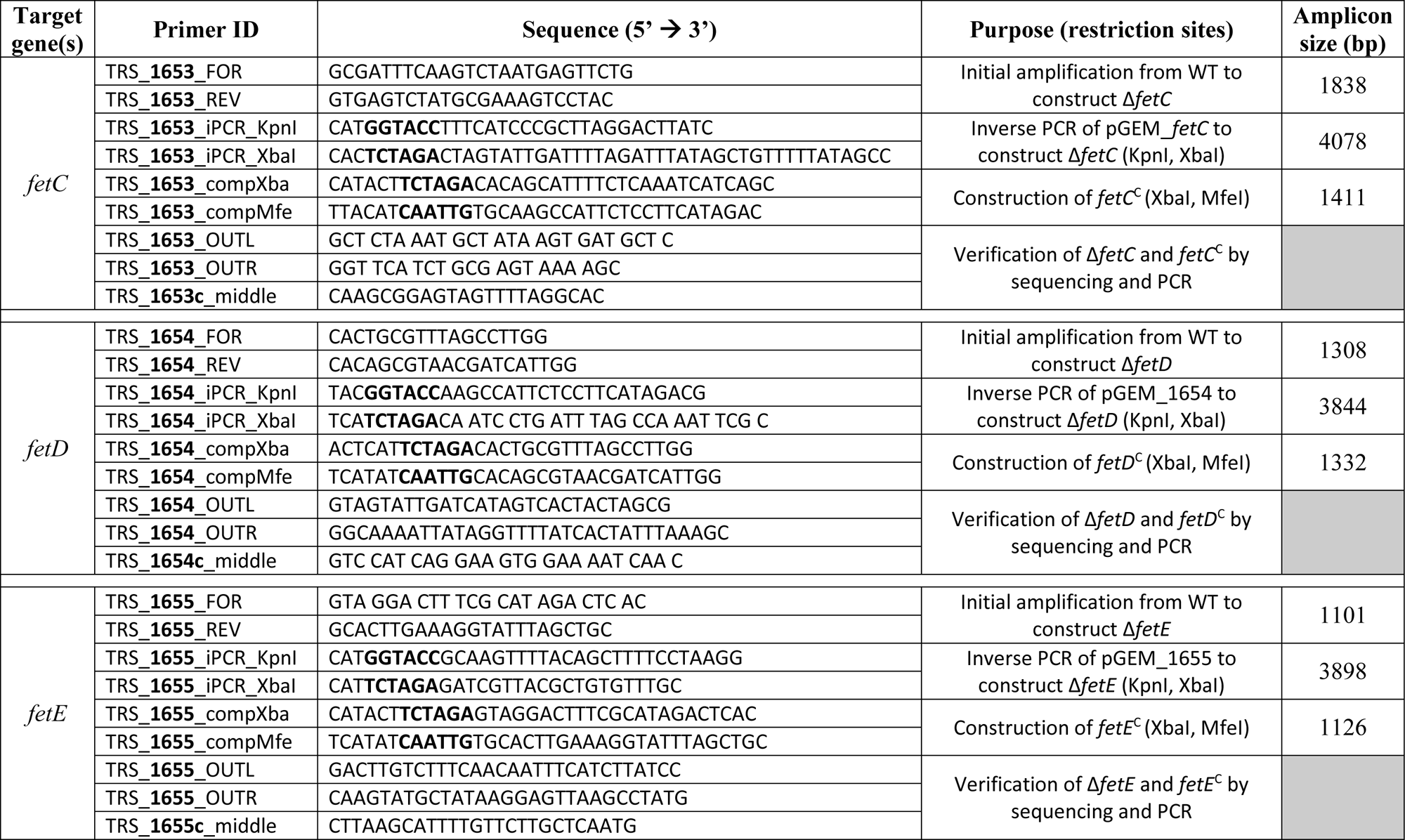
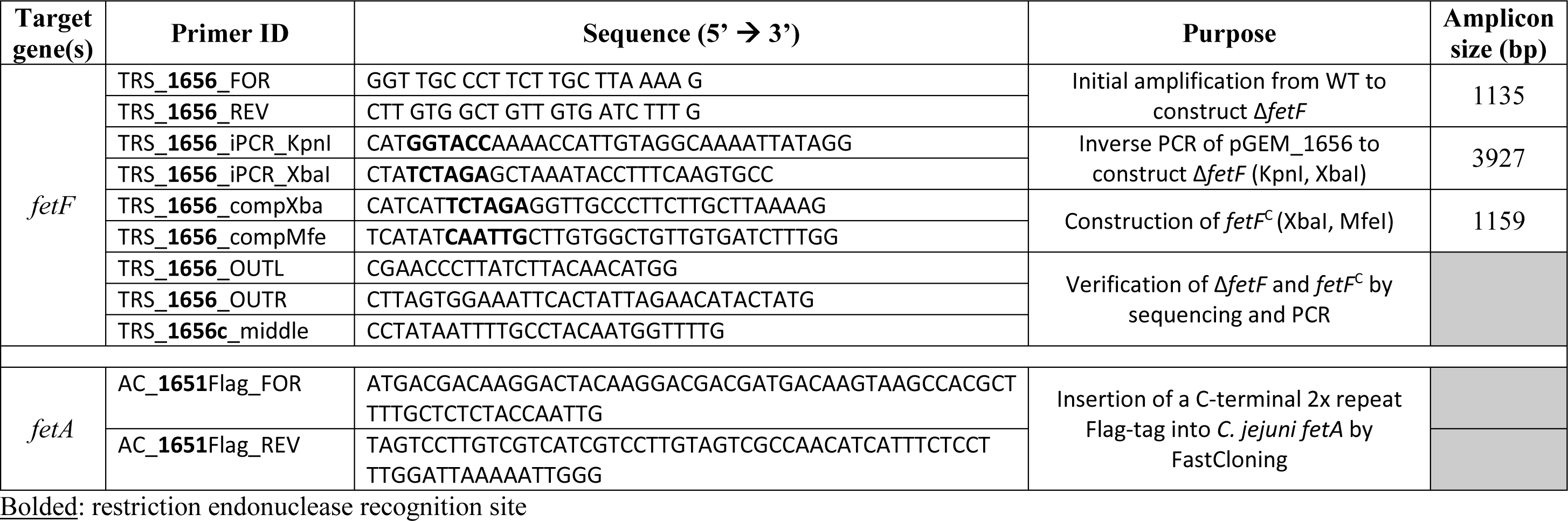
Bacterial strains, plasmids, and primers used in this study.

**Supplemental Table S2:** Data collection and refinement statistics for FetE.

| Data Collection<sup>a</sup> | FetE (iodide) | FetE (native) |
| --- | --- | --- |
| Resolution Range (Å) | 36.66-1.94 (1.99-1.94) | 46.73-1.50 (1.60-1.50) |
| Space group | $P2_1$ | $P2_1$ |
| Cell dimensions (Å) | $a = 49.22$ $b = 43.38$ $c = 72.20$ | $a = 49.25$ $b = 43.38$ $c = 71.86$ |
| | $\beta = 108.3$ | $\beta = 108.4$ |
| Wavelength (Å) | 1.600 | 0.979 |
| Unique Reflections | 20879 | 46005 |
| Completeness (%) [Anom] | 97.1 (86.0) [94.7 (75.8)] | 99.2 (96.9) |
| Average I/σI | 13.7 (4.3) | 16.7 (4.1) |
| Redundancy [Anom] | 4.3 [2.2] | 4.2 |
| $R_{\text{meas}}$ | 0.089 (0.286) | 0.052 (0.267) |
| $CC_{1/2}$ | 0.995 (0.941) | 0.999 (0.935) |
| **Refinement** |  |  |
| $R_{\text{work}} / R_{\text{free}}$ | | 0.156 / 0.183 |
| No. of waters |  | 272 |
| Average $B$-value (Å<sup>2</sup>) | | 28.5 |
| r.m.s.d. bond lengths (Å) |  | 0.009 |
| Ramachandran plot |  |  |
| Most-favorable (%) |  | 96.8 |
| Allowed (%) |  | 3.2 |
| Outliers (%) |  | 0 |

<sup>a</sup> Values for the highest resolution shell are shown in parenthesis

